# Convergent lignocellulose degradation in terrestrial crabs is driven by distinct host genus-specific microbial communities

**DOI:** 10.64898/2026.04.24.720673

**Authors:** Victoria M. Watson-Zink, Laetitia G.E. Wilkins, Jonathan A. Eisen, Richard K. Grosberg, Alexandra Hiller, Cassandra L. Ettinger

**Affiliations:** Department of Organismic and Evolutionary Biology, Harvard University, Cambridge, Massachusetts, United States; Amt für Jagd und Fischerei Graubünden, Abteilung Fischerei, Ringstrasse 10, 7000 Chur, Switzerland; Department of Evolution and Ecology, University of California, Davis, United States 95616; Smithsonian Tropical Research Institute, Apartado 0843-03092, Balboa, Ancon, Republic of Panamá; Department of Microbiology and Plant Pathology, University of California, Riverside, Riverside, California, United States

**Keywords:** land crabs, sea-to-land transitions, metagenomics, gut microbiome, metagenome-assembled genomes, lignocellulose degradation, phylosymbiosis, Anomura, Brachyura

## Abstract

**Background:** Restrictions on the types of food available on land have repeatedly triggered the convergent evolution of herbivory in terrestrial animals. This pattern also holds true in many terrestrially adapted crabs, which have independently colonized land more than 17 times since the Cretaceous. Many clades of terrestrial crabs are now almost completely herbivorous, standing in contrast to the ancestral pattern of detrivory. While many bacteria possess efficient pathways for degrading lignin and cellulose, the role of gut microbiomes in facilitating these dietary shifts in terrestrial crabs remains poorly understood. To explore the relationship between microbial community structure and the ability of land crabs to digest lignocellulose, we conducted read-based and assembly-based metagenomic analyses on feces collected from the guts of 14 crab species across six genera, representing a gradient of terrestriality from the lower intertidal zone to forested habitats.

**Results:** We generated 129 metagenome-assembled genomes (MAGs) that represent key members of these gut microbial communities, establishing a foundational resource for future studies on crab-microbiome interactions. We found that host genus explained most of the variation in bacterial community composition, while degree of terrestrial adaptation (i.e. terrestrial grade) explained a smaller proportion. We also identified multiple bacterial genera that strikingly differed in relative abundance across terrestrial grades, crab genera, and diet type. Broad-scale functional analyses of general carbon metabolism across crabs revealed an absence of complete pathways in crabs from lower terrestrial grades, suggesting a functional divergence in gut communities linked to habitat transition. Fine-scale functional analyses of predicted carbohydrate-active enzyme (CAZyme) domains allowed us to connect specific MAGs to lignocellulose degradation pathway genes, demonstrating that different crab genera harbor distinct microbial taxa that may have similar CAZyme profiles in their guts.

**Conclusions:** This work provides a foundational metagenomic resource for genomic exploration of microbial communities in terrestrial crab guts. These results suggest that the gut microbiomes of terrestrially adapted crabs are structured primarily by host identity and may have convergently acquired microbes with similar functions to help perform lignocellulose degradation. Overall, different degrees of adaptation to terrestrial environments, including resulting dietary shifts, may be responsible for functional divergence in crab gut community assembly.

## Background

All life began in the oceans but most macroscopic species on Earth currently reside on land [1]. Sea-to-land transitions are rare across evolutionary time however, suggesting that these transitions present major challenges to nearly every critical physiological and biological process, including reproduction, development, excretion, respiration, osmoregulation, sensory reception, and nutrient acquisition [2, 3]. In particular, restrictions on the types of food available on land has repeatedly triggered the evolution of herbivory in metazoan lineages across the Tree of Life, as seen in terrestrial vertebrates [4], insects [5], isopods [6] and blennies [7]. The reverse has also been documented: as cetaceans recolonized the oceans, there was a concurrent shift from herbivory to carnivory [8], suggesting that dietary shifts play a key role in the colonization of novel habitats.

Decapod crabs are an exceptional example of an ancestrally marine clade that has recently and repeatedly colonized terrestrial environments, representing one of the most dramatic evolutionary stories in extant marine organisms. Since the mid-Cretaceous, there have been more than seventeen convergent transitions to land in the decapod infraorder Brachyura (containing the “true crabs”) alone [9], as well as a single independent colonization in Anomura by the terrestrial hermit crabs in Coenobitidae. Compared to their marine counterparts that consume mostly animal prey, micro- and macroalgae, and detritus [10], land crabs have adapted to extracting nutrients from a completely different suite of items that grow abundantly in the terrestrial realm: land plants. The diet of *Gecarcoidea natalis*, the Christmas Island red crab, for example, contains approximately 97% plant matter (leaf litter: 71%, fruits: 21%, wood 3%, and flowers: 2%) [11], making this crab effectively herbivorous [12]. But land plants possess many complex structural polymers that make them difficult to digest, with lignin and lignocellulose chief among these [13]. Few, if any, invertebrates can digest these materials endogenously [14]. Here we ask whether the repeated formation of beneficial symbioses between proto-land crabs and lignin-digesting microbes, and the resulting ability to exploit a novel and abundant food source, may have been a key event facilitating the terrestrialization of land crabs.

A recent study suggested that the gut microbiomes of terrestrial crabs may have facilitated this dramatic shift in diet [15]. Populations of the same species of terrestrial hermit crab grew to different sizes on their respective islands based solely on the relative availability of seagrass debris, which is an easily digestible food. But when fed a diet of dicot leaves, which are a low quality, recalcitrant food source that grows abundantly on the islands lacking seagrass debris, local crabs that were “acclimated” to eating dicot leaves grew faster than the crabs acclimated to eating only seagrass debris. One untested explanation was that the crab populations host different microbial communities, leading to the observed differences in foodtype acclimation and growth.

While recent work has investigated the importance and putative functions of beneficial digestive microbiomes in intertidal and mangrove crabs [16, 17], no studies published to date have explicitly and directly examined the gut microbiomes of crabs with greater degrees of terrestrial adaptation, or how this partnership affects a crabs’ ability to subsist almost entirely on lignified plant material (see Cannicci et al. [18] for a review promoting the study of intertidal and terrestrial crabs as an ideal model system for exploring the role of gut microbiota in the repeated evolutionary leap from sea to land).

In this study, we explored the relationship between land crab gut microbial assembly and the ability to digest recalcitrant plant matter, predicting that land crabs that display greater degrees of terrestrial adaptation will possess microbes in their guts that are putatively capable of digesting recalcitrant plant material (i.e. lignin, lignocellulose, hemicellulose and cellulose). We sampled the fecal microbiota of several species of land crabs that display differing degrees of terrestrial adaptation and sought to understand whether there were microbes present in their guts that might play a key role in lignocellulose degradation. Looking across the gradient of terrestriality in crabs (described by Watson-Zink [2] as a series of terrestrial “grades” from I to VI, with subtidal species existing at lowest grades and arid-zone dwelling species at the highest grades), we also aimed to determine whether there is a relationship between relative terrestrial grade and microbial community composition.

This study will add to the growing literature that evaluates the role of the microbiome as an integral part of our definition of what an organism is in the context of a major ecological transition. Our findings will also better characterize the ecological contexts that may have enabled the many transitions onto land by the decapod crabs, and how the formation of beneficial mutualistic microbial relationships may have paved the way for many of these species to adapt to novel terrestrial environments.

## Methods

### Field collections

We hand-collected intermolt adult males of *Tuerkayana celeste* (n=4), *Gecarcoidea natalis* (n=4), *Birgus latro* (n=3), *Geograpsus grayi* (n=1), and *Ocypode ceratophthalmus* (n=1) from Christmas Island National Park in Australia and obtained intermolt adult males of *T. magna* (n=4) and *G. humei* (n=1) from the pet trade in Singapore (but the animals were originally collected in Java, Indonesia) (Figure 1, Table 1). Since many of the Australian species are protected and reside in a national park, we obtained research, collection, and export permits from the Department of the Environment and Energy of the Australian Government (Permit Numbers: PWS2017-AU-001942, AU-COM2017-354, AU-COM2017-363, and AU-COM2017-373) to complete this study. We confirmed species by gross morphology and, in the case of *T. magna*, also via gonopod morphology. Once in the laboratory, we kept all crabs in individual 10 L plastic buckets at ambient air temperatures (which ranged from 28 °C - 41 °C) and humidity (which ranged from 60% - 82%). We filled each bucket with 1.5 L Christmas Island freshwater (CIFW), which we either collected from a small stream at Ross Hill Gardens, Christmas Island, Australia (10°29.0923’S, 105°40.8031’E) for sampling on Christmas Island, or mixed ourselves following the methods of Wood et al. (1986) for sampling in Singapore where natural CIFW was not available. We refreshed the water in each bucket every 12 h until dissection.

**Figure 1.**
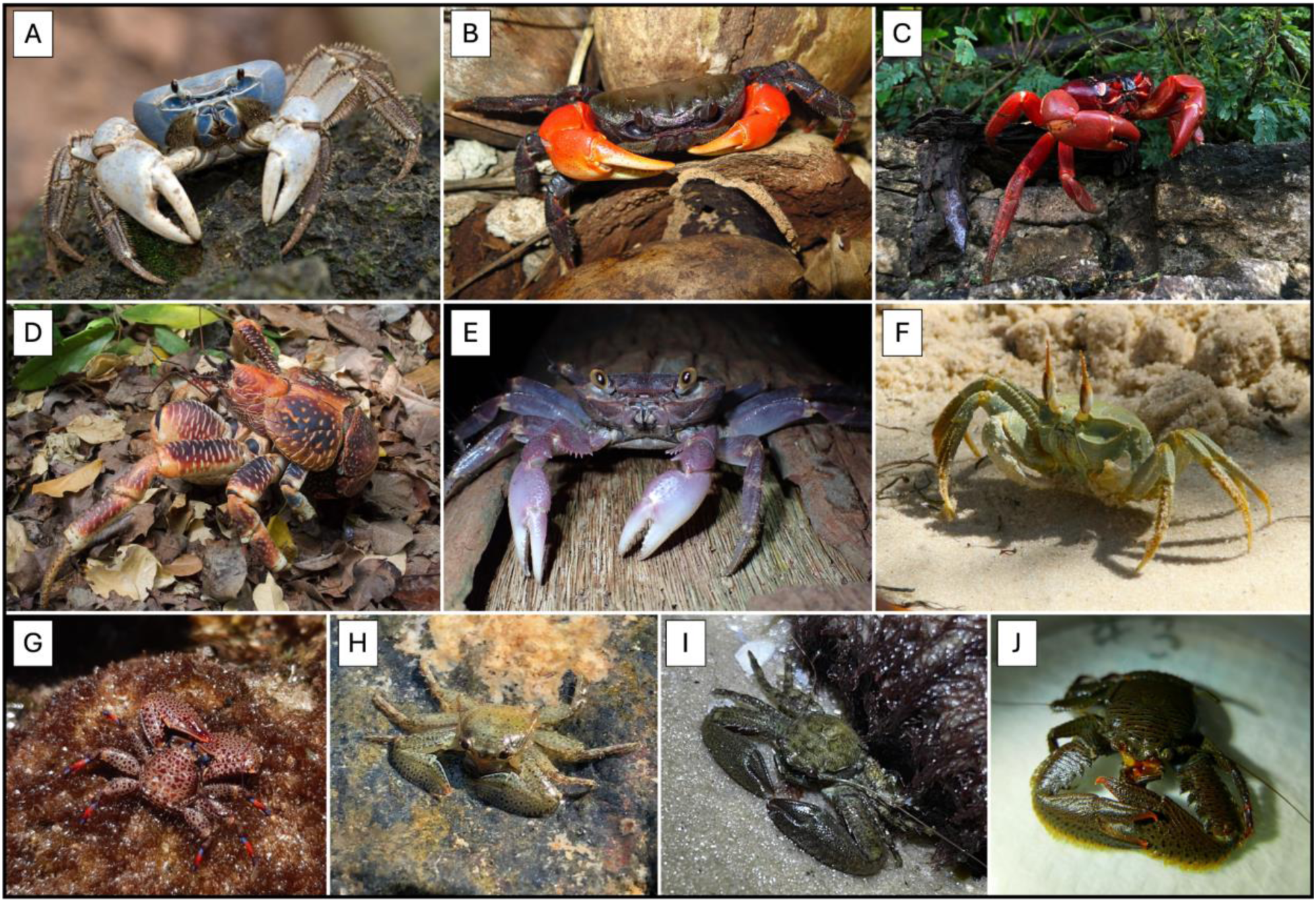
Representative species used in this study. (A) Tuerkayana celeste; (B) Tuerkayana magna; (C) Gecarcoidea natalis; (D) Birgus latro; (E) Geograpsus grayi; (F) Ocypode ceratophthalmus; (G) Petrolisthes donadio; (H) Petrolisthes nobilii; (I) Petrolisthes armatus; (J) Petrolisthes sp. All images are licensed under CC BY-NC 4.0 by budak, dj_maple, danicalockett, Watson-Zink, V.W., adurbano, Leggatt, N., carisa, puravida_ac, cowturtle, and Wilkins, L.G.E., respectively.

**Table 1.** Summary of crabs collected for use in this study. Here we provide details on each metagenomic sample used in this study including host crab taxonomic information, collection location, collection year, terrestrial grade, diet, and the NCBI accession number for the metagenomic reads.

We collected porcelain crabs by hand in the intertidal zone in Panamá with collection permission from the Ministerio de Medio Ambiente de Panamá (Permit Number SE/A-111-17 and SE/A-72-18). We transported individuals to the laboratory in 50 mL Falcon tubes with 35 mL of seawater from their natural habitat, where we held them in 10 L plastic buckets at ambient air temperatures for 24 h before dissection. Porcelain crabs included 12 *Petrolisthes* species that were identified to the species level by morphology (Table 1): *P. armatus* (n = 2; sampled at the Smithsonian Tropical Research Institute “STRI” in Bocas del Toro; mangrove habitat in the Caribbean Sea), *P. bolivarensis* (n = 1; STRI in Bocas del Toro; coral habitat in the Caribbean Sea), *Petrolisthes coeruleus* (n = 2; STRI in Bocas del Toro; coral habitat in the Caribbean Sea), *P. armatus* (n = 2; Playa Hermosa; rocky habitat in the Pacific Ocean), *P. donadio* (n = 3; Islas Secas; coral habitat in the Pacific Ocean), *P. haigae* (n = 1; Islas Secas; coral habitat in the Pacific Ocean), *P. nobilii* (n = 1; Islas Secas; coral habitat in the Pacific Ocean).

### Molecular methods and sequence generation

In the field using tools sterilized with 100% ethanol, we dissected out the entire digestive tracts of the larger crabs (i.e., crabs in the genera *Tuerkayana*, *Ocypode*, *Geograpsus*, *Gecarcoidea*, and *Birgus latro*). We stored all field samples in microcentrifuge tubes containing 1 mL Zymo DNA/RNA Shield (Zymo Research, R1100-250) (or 5 mL for *Birgus latro*, which had a digestive tract over 0.25m long), and flash froze them first in liquid nitrogen and then transported them back to the United States in a dewar on dry ice pellets. We then stored them at -80C until further processing. In the laboratory using 100% ethanol flame-sterilized tools, we scraped any visible fecal matter from the digestive tracts of the larger crabs while deliberately avoiding sampling host tissue, but dissected out the entire gut from *Petrolisthes sp*. due to their much smaller size.

We extracted DNA from the samples using a Zymo Research DNA Mini Kit following the manufacturer’s instructions with two modifications: During Step 2, we bead beat the samples for a total of 5 minutes split across two 2.5 minute sessions with a brief rest to prevent overheating, and during Step 13, we centrifuged the samples for 3 minutes. We quantified DNA concentrations for each sample using a Qubit 4 Fluorometer (Qiagen). We also extracted DNA from a no-sample added kit control (i.e., DNA extracted from kit reagents) for use as a negative control during analyses.

We checked the DNA quality of each sample on a 1% E-Gel. In total, we provided 33 DNA samples (32 fecal samples and 1 negative kit control) to the UC Davis Genome Center DNA Technologies Core (https://dnatech.genomecenter.ucdavis.edu/) for library preparation. They were sequenced on an Illumina NovaSeq Sp500 as paired-end 250 bp reads. We deposited the metagenomic sequence reads generated for this project in NCBI’s GenBank (BioProject ID [PRJNA1141481](http://www.ncbi.nlm.nih.gov/bioproject/1141481)).

### Host mitogenome phylogenetics

We passed raw reads to MitoZ (v3.4) for mitogenome assembly and mitochondrial protein coding gene identification and annotation [19]. We used the GenBank files (.gbf) that are included in the MitoZ output and the complete mitogenome of *Upogebia bowerbankii* as an outgroup (NCBI accession number NC_041158.1) to assemble a host phylogeny using MitoPhAST (v3.0) [20] (Figure 2).

**Figure 2.**
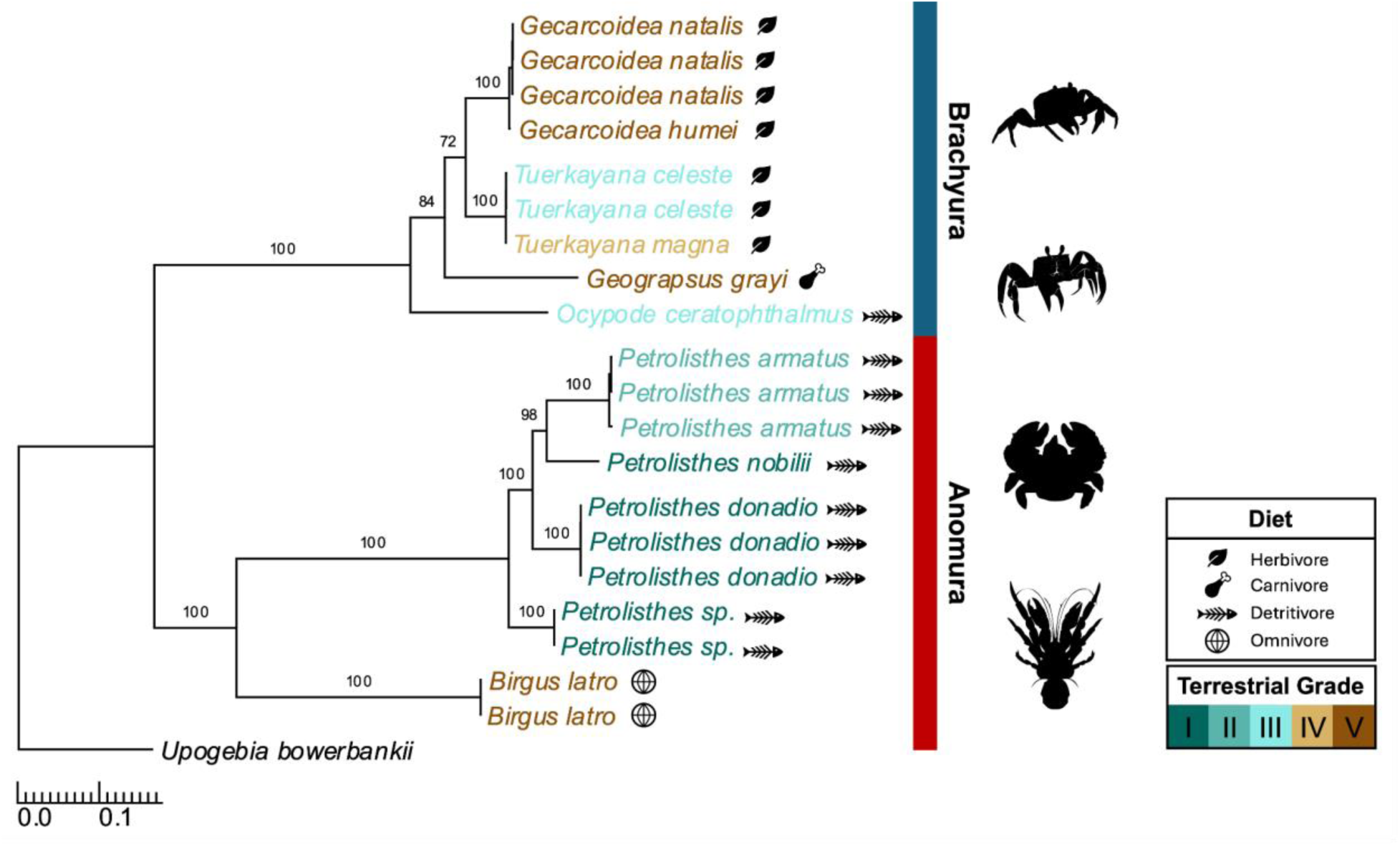
Mitogenome maximum likelihood host phylogeny with bootstrap branch support. Species name colors reflect respective terrestrial grades from I to V. Adjacent icons reflect diet type (i.e., herbivore, carnivore, detritivore, or omnivore) and colored horizontal bars distinguish the two decapod crab infraorders. Silhouette images provided by Joanna Wolfe (*Lomis hirta*), Loran Honório da Silva (*Ocypode quadrata*), Margot Michaud (*Gecarcinus quadratus*), and T. Michael Keesey (*Birgus latro*).

### Read-based taxonomic profiling

We trimmed, PhiX de-contaminated, and quality checked our raw Illumina reads using BBMAP v37.61’s BBDUK feature [21]. We trimmed sequences by q=15 and dropped sequences that were shorter than 149 bp. We taxonomically classified the trimmed reads using Kraken2 v. 2.1.2 [22] against the NCBI non-redundant nucleotide database (downloaded June 16, 2022). We then ran Bracken v.2.5 [23], which uses a Bayesian re-estimation method to better assign reads based on genome lengths from the reference database, on the Kraken2 outputs to estimate the number of reads associated with each species in each individual sample. We used ETE3 [24] to obtain the full taxonomic lineage string of each identified species from NCBI.

### Metagenomic assembly, annotation and binning

We combined clean read libraries from all crabs into co-assemblies using MEGAHIT v. 1.2.9 [25]. We binned the resulting metagenomic assembly scaffolds using a combination of MetaBAT2 v. 2.15 [26] with and without read coverage information, MaxBin2 v. 2.2.7 [27], BinSanity v. 0.5.4 [28], and CONCOCT v1.1.0 [29]. To improve overall binning performance given that some samples contained host crab DNA, we also generated three sub-sampled co-assemblies: one for all porcelain crabs (Detritivores), one for *Tuerkayana* and *Gecarcoidea* (Herbivores), and one for *Birgus*, *Ocypode* and *Geograpsus* (Omnivores/Carnivores). We passed these sub-sampled co-assemblies to WhoKaryote v. 1.1.2 [30] which we used to identify bacterial contigs. We then binned the resulting bacterial contigs as described above.

We optimized draft genome bins (metagenome-assembled genomes ‘MAGs’) from the entire co-assembly and the sub-sampled co-assemblies and we created a non-redundant set using DAS Tool v. 1.1.6 [31]. We checked optimized MAGs for completion using CheckM2 v. 1.0.1 [32]. We further refined bins using both charcoal [33] and MAGpurify [34] to detect and remove contamination. We referred to MAGs that were determined to be 90% or more complete and less than 10% contaminated post refinement as high-quality MAGs [35]. We used GTDB-Tk v.2.2.6 [36] to assign putative taxonomies to MAGs using a combination average nucleotide identity (ANI) and phylogenetic placement. We deposited the final MAG set generated for this project in NCBI’s GenBank (BioProject ID [PRJNA1141481](http://www.ncbi.nlm.nih.gov/bioproject/1141481)).

In order to investigate the relative abundance of MAGs across samples, we used coverM v. 0.6.1 [37] with minimap2 v. 2.24 [38] to calculate RPKM (reads per kilobase per million reads mapped) with the following parameters: --min-read-aligned-percent 0.75 --min-read-percent-identity 0.95 --min-covered-fraction 0. We chose these parameters to capture rare reads, as we expected some samples to have low coverage due to host DNA, while increasing specificity thresholds to reduce spurious mapping.

### Taxonomic composition analysis

We imported the count tables from read-based analysis using Kraken2/Bracken (hereafter K2/B) and MAG-based analyses using coverM into R v. 4.3.0 for downstream analysis and visualization using the phyloseq v. 1.44.0, tidyverse v. 2.0.0 and vegan v. 2.6.6.1 packages [39–42]. We separated K2/B data into three datasets: one each for bacteria, archaea, and fungi.

We used decontam’s prevalence method to identify possible contaminants with a higher prevalence in the negative control than in true samples and ran decontam with a threshold of 0.5 [43]. We then removed all predicted contaminants and the negative control sample before downstream analysis.

To assess differences in beta diversity, we calculated Bray Curtis dissimilarities using the distance function from phyloseq and visualized them using a principal coordinate analysis (PCoA) plot with the ordinate function in phyloseq. We then tested for significant differences in beta diversity centroids (i.e., means of each group of interest) across crab genera (composed of 9 *Tuerkayana* samples, 5 *Gecarcoidea* samples, 3 *Birgu*s samples, 12 *Petrolisthes* samples, 1 *Geograpsus* sample, and 1 *Ocypode* sample), terrestrial grades (composed of 7 samples for Grade I, 5 samples for Grade II, 5 samples for Grade 3, 4 samples for Grade 4, and 11 samples for Grade V), and diet (represented by 13 herbivore samples, 13 detritivore samples, or 5 omnivore samples and 1 carnivore sample) using permutational multivariate analyses of variance (PERMANOVAs) with 9,999 permutations and by = “margin” and by = “sequential” with shuffled terms using the adonis2 function. Ocean basin was included as a covariate in the PERMANOVA models. In sequential models where crab genus was placed before diet, diet was dropped from the model and crab genus was able to explain the variation otherwise attributed to diet. When diet was placed before crab genus, both terms were maintained with variation split between them. We interpreted this to mean that crab genus encompasses the variation due to diet, as well as additional variation. Thus, we did not include diet in the final model choice and the final model was only used to assess the marginal effects of crab genus and terrestrial grade. The final model was Genus + Terrestrial grade + Ocean with by= “margin”.

We used *post hoc* pairwise PERMANOVAs to test pairwise differences using the pairwise.adonis function from the pairwiseAdonis package [44]. We also assessed differences in dispersion (i.e., variability) using the betadisper and permutest functions from the vegan package with 9,999 permutations. For tests that resulted in a rejected null hypothesis, we used *post hoc* Tukey’s Honest Significant Difference (HSD) tests to determine which pairwise differences might drive the significant dispersion differences. We corrected all *p*-values using the Benjamini-Hochberg method.

To assess differences in the relative abundance of microbial genera, we first transformed counts into proportions. For statistical analysis, we filtered out low abundance taxa representing less than 1% mean abundance across each dataset. We then tested for significant differences in the relative abundance of microbial genera across host genera, terrestrial grades (I-V), and diet (herbivore, detritivore or omnivore/carnivore) using Kruskal-Wallis (K-W) tests with 9,999 permutations. For tests that resulted in a rejected null hypothesis, we performed *post hoc* Dunn tests to assess driving pairwise contrasts. We corrected all *p*-values using the Benjamini-Hochberg method.

### Functional analysis

We annotated MAGs using the Distilling and Refining Annotations of Metabolism (DRAM) v. 1.5.0 workflow [45]. Briefly, DRAM predicts genes using Prodigal [46] and then annotates these genes using searches against the KOfam [47], PFAM [48], dbCAN v. 11 [49, 50], and MEROPS [51] databases. DRAM retains a dbCAN HMM hit only if its sequence E-value is ≤ 1×10⁻¹⁸ and the HMM coverage is ≥ 35%, which is stricter than the dbCAN default (E ≤ 1×10⁻¹⁵). We extracted all genes with predicted CAZyme (Carbohydrate Active Enzymes) family assignments matching families reported in the literature to be related to lignocellulose degradation [6, 52–57]. To reduce spurious calls, we excluded genes that Prodigal flagged as partial (truncated at a contig edge). Because DRAM reports overlapping family and subfamily HMM hits for the same protein region, we counted each CAZy family once per gene (gene-family hits) rather than counting individual HMM hits. We then visualized the detection frequency of gene-family hits across MAGs using the tidyverse v. 2.0.0 [40] and pheatmap v. 1.0.12 [58] in R v. 4.3.0 [39]. For visualization purposes, CAZyme gene-family frequencies were log-transformed.

We assembled individual metagenomic samples using spades v. 3.13.1 [59] and then used WhoKaryote v. 1.1.2 [30] to identify bacterial contigs. We then annotated the bacterial contigs from each metagenomic assembly with METABOLIC v.4.0 using ‘METABOLIC-G.pl’ [60]. We visualized heatmaps of detected functional genes and pathways using the tidyverse v. 2.0.0 [40] in R v. 4.3.0 [39]. Due to the smaller size of our dataset, we implemented a random forest model using ranger v. 0.16.0 [61] and caret v. 6.0.94 [62] to preliminarily explore whether the count of genes in complex carbon degradation pathways could accurately be used to predict crab diet. We split the dataset into 70% training and 30% testing subsets, maintaining the proportional distribution of diet groups via stratified sampling. We used class weights to handle the sample imbalance between dietary groups by giving Carnivore/Omnivore higher importance (x2). Due to the small nature of the dataset, cross-validation was performed 10-fold, repeated 10 times. The optimal model parameters were determined based on the highest accuracy. The final Random Forest model was trained with 500 trees, using 6 predictors (mtry=6), with the Gini split rule. Feature importance was calculated using permutation importance.

## Results

Although we intentionally attempted to avoid sampling and sequencing host tissue, we nonetheless were able to successfully assemble complete crab mitogenomes from 24 of our 33 samples; 14 from Anomura and 10 from Brachyura. The mitogenomes varied in size from 14,817 bp (*Tuerkayana magna*) to 18,276 bp (*Birgus latro*). When placed into a phylogeny (Figure 2), the mitogenome sequences reflected phylogenetic relationships within each infraorder that are consistent with existing literature (Wolfe et al. 2024).

### Comprehensive high-quality bacterial MAG collection

In total, we assembled 129 MAGs with >90% completion including 114 high-quality MAGs (< 5% contamination) and 15 medium-quality MAGs (< 10% contamination) (Table 2). Due to changes in taxonomic naming conventions and systematics, each mention of a microbial clade will be followed in parenthesis with that clade’s previous name where applicable. Most of these MAGs belong to the following clades: Bacteroidota (i.e., Bacteroidetes: 35), Pseudomonadota (i.e., Proteobacteria: 34), Bacillota (i.e., Firmicutes: 30) and Actinomycetia (i.e., Actinobacteriota: 18). Despite high completion levels in the MAG collection, GTBD-Tk was only able to place some MAGs at higher taxonomic levels (i.e., class, order, family), likely as a result of evolutionary novelty relative to the GTDB database (Table S1).

**Table 2.** Summary of genomic features of 129 draft MAGs. Genomic features are summarized for each MAG, including putative taxonomic identity in both the GTDB and NCBI databases, MAG size (bp), N50, GC content, number of genes, completion and contamination estimates as generated by CheckM2, and NCBI accession number. MAGs are sorted by CheckM2 completion. See Table S1 for full GTDB taxonomy.

| **Bin ID** | **GTDB Taxonomy** | **NCBI Taxonomy** | **Completeness** | **Contamination** | **MAG size (bp)** | **N50 (bp)** | **GC (%)** | **No. of genes** | **NCBI accession no.** |
|---|---|---|---|---|---|---|---|---|---|
| VLC_001 | CAG-239 *sp.* | *Alphaproteobacteria bacterium* | 100 | 0.36 | 1812003 | 142053 | 0.34 | 1865 | JBKDZH000000000 |
| VLC_002 | *Ottowia sp.* | *Ottowia sp.* | 100 | 1.32 | 3190115 | 54528 | 0.62 | 3015 | JBKDZI000000000 |
| VLC_003 | *Demequina sp.* | *Demequina sp.* | 100 | 1.75 | 2855939 | 117555 | 0.68 | 2661 | JBKDZJ000000000 |
| VLC_004 | JACXVD01 *sp.* | *Sphingomonadaceae bacterium* | 100 | 3.47 | 3202755 | 14049 | 0.55 | 3340 | JBKDZK000000000 |
| VLC_005 | *Niabella sp.* | *Niabella sp.* | 100 | 0.15 | 3923749 | 43839 | 0.37 | 3447 | JBKDZL000000000 |
| VLC_006 | *Denitromonas sp.* | *Denitromonas sp.* | 100 | 2.49 | 4311116 | 64139 | 0.66 | 4084 | JBKDZM000000000 |
| VLC_007 | *Brachymonas sp.* | *Brachymonas sp.* | 100 | 0.02 | 2020663 | 86269 | 0.61 | 1884 | JBKDZN000000000 |
| VLC_008 | *Malaciobacter mytili* | *Malaciobacter mytili* | 100 | 0.35 | 2761445 | 102896 | 0.26 | 2783 | JBKDZO000000000 |
| VLC_009 | *Phascolarctobacterium faecium* | *Phascolarctobacterium faecium* | 100 | 0.22 | 2339828 | 133075 | 0.44 | 2154 | JBKDZP000000000 |
| VLC_010 | UBA10566 sp002399565 | *Bacteroidales bacterium* | 100 | 1.16 | 2451276 | 37027 | 0.31 | 2195 | JBKDZQ000000000 |
| VLC_011 | *Fusobacterium_A ulcerans* | *Fusobacterium ulcerans* | 99.99 | 0.76 | 3545176 | 30312 | 0.3 | 3454 | JBKDZR000000000 |
| VLC_012 | *Parabacteroides gordonii* | *Parabacteroides gordonii* | 99.99 | 3.77 | 6691659 | 46561 | 0.44 | 5343 | JBKDZS000000000 |
| VLC_013 | *Enterococcus_B sp.* | *Enterococcus sp.* | 99.97 | 3.1 | 2462183 | 36207 | 0.37 | 2413 | JBKDZT000000000 |
| VLC_014 | *Flavobacteriaceae sp.* | *Flavobacteriaceae bacterium* | 99.96 | 1.83 | 3292456 | 60231 | 0.41 | 2870 | JBKDZU000000000 |
| VLC_015 | *Bacteroides reticulotermitis* | *Bacteroides reticulotermitis* | 99.96 | 2.64 | 5427199 | 74197 | 0.43 | 4309 | JBKDZV000000000 |
| VLC_016 | *Xanthomarina gelatinilytica* | *Xanthomarina gelatinilytica* | 99.94 | 0.02 | 3259468 | 44161 | 0.35 | 3107 | JBKDZW000000000 |
| VLC_017 | *Vibrio sp.* | *Vibrio sp.* | 99.9 | 0.7 | 3648598 | 89984 | 0.41 | 3281 | JBKDZX000000000 |
| VLC_018 | *Bacteroides sp.* | *Bacteroides sp.* | 99.89 | 0.82 | 3212730 | 54058 | 0.41 | 2627 | JBKDZY000000000 |
| VLC_019 | *Draconibacterium sp.* | *Draconibacterium sp.* | 99.85 | 0.37 | 5363764 | 25278 | 0.41 | 4468 | JBKDZZ000000000 |
| VLC_020 | *Vibrio fluvialis* | *Vibrio fluvialis* | 99.85 | 9.24 | 4960588 | 15481 | 0.51 | 5133 | JBKEAA000000000 |
| VLC_021 | *Enterococcus_C sp.* | *Enterococcus sp.* | 99.84 | 0.71 | 2093463 | 51225 | 0.4 | 2115 | JBKEAB000000000 |
| VLC_022 | *Citrobacter portucalensis* | *Citrobacter portucalensis* | 99.82 | 0.54 | 3877956 | 15342 | 0.53 | 3951 | JBKEAC000000000 |
| VLC_023 | *Shewanella sp.* | *Shewanella sp.* | 99.81 | 2.74 | 4653781 | 76724 | 0.47 | 4139 | JBKEAD000000000 |
| VLC_024 | *Limnobaculum xujianqingii* | *Limnobaculum xujianqingii* | 99.77 | 0.07 | 3572954 | 35203 | 0.45 | 3263 | JBKEAE000000000 |
| VLC_025 | *Aestuariibaculum sp.* | *Aestuariibaculum sp.* | 99.55 | 0.01 | 2826470 | 22158 | 0.36 | 2506 | JBKEAF000000000 |
| VLC_026 | UBA660 *sp.* | *Bacilli bacterium* | 99.51 | 0.44 | 1531152 | 38930 | 0.29 | 1581 | JBKEAG000000000 |
| VLC_027 | *Pseudorhodobacter sp.* | *Pseudorhodobacter sp.* | 99.48 | 0.69 | 3464994 | 66558 | 0.59 | 3340 | JBKEAH000000000 |
| VLC_028 | UJ101 sp. | *Flavobacteriales bacterium* | 99.35 | 0.05 | 2287072 | 81435 | 0.33 | 2280 | JBKEAI000000000 |
| VLC_029 | *Jhaorihella sp.* | *Jhaorihella sp.* | 99.12 | 4.17 | 3859280 | 64995 | 0.65 | 3789 | JBKEAJ000000000 |
| VLC_030 | *Mangrovibacterium sp.* | *Mangrovibacterium sp.* | 99.02 | 0.02 | 2568125 | 79325 | 0.38 | 2141 | JBKEAK000000000 |
| VLC_031 | *Cumulibacter sp.* | *Cumulibacter sp.* | 98.77 | 4.26 | 3923282 | 66537 | 0.63 | 3823 | JBKEAL000000000 |
| VLC_032 | *Dysgonomonas sp.* | *Dysgonomonas sp.* | 98.66 | 7.88 | 4279992 | 23723 | 0.37 | 4088 | JBKEAM000000000 |
| VLC_033 | *Mycobacterium sp.* | *Mycobacterium sp.* | 98.65 | 1.7 | 3372817 | 41632 | 0.67 | 3188 | JBKEAN000000000 |
| VLC_034 | *Lachnospiraceae sp.* | *Lachnospiraceae bacterium* | 98.62 | 0.76 | 2978259 | 58811 | 0.45 | 2753 | JBKEAO000000000 |
| VLC_035 | UBA10566 sp. | *Bacteroidales bacterium* | 98.46 | 3.65 | 2615557 | 22950 | 0.32 | 2545 | JBKEAP000000000 |
| VLC_036 | *Bacteroidales sp.* | *Bacteroidales bacterium* | 98.42 | 2.27 | 2581534 | 36828 | 0.4 | 2212 | JBKEAQ000000000 |
| VLC_037 | *Fimivivens sp.* | *Fimivivens sp.* | 98.34 | 0.16 | 1977360 | 243125 | 0.49 | 1896 | JBKEAR000000000 |
| VLC_038 | *Marinicella pacifica* | *Marinicella pacifica* | 98.13 | 0.1 | 2871122 | 399837 | 0.45 | 2579 | JBKEAS000000000 |
| VLC_039 | *Anaerorhabdus sp.* | *Anaerorhabdus sp.* | 98.03 | 0 | 1441107 | 50830 | 0.35 | 1430 | JBKEAT000000000 |
| VLC_040 | *Tetrasphaera sp.* | *Tetrasphaera sp.* | 97.92 | 0.76 | 3033711 | 55004 | 0.67 | 2945 | JBKEAU000000000 |
| VLC_041 | *Anaerorhabdus sp.* | *Anaerorhabdus sp.* | 97.73 | 2.58 | 2639509 | 55137 | 0.33 | 2828 | JBKEAV000000000 |
| VLC_042 | *Aminipila sp.* | *Aminipila sp.* | 97.72 | 0.44 | 3035754 | 48364 | 0.35 | 2916 | JBKEAW000000000 |
| VLC_043 | *Parahaliea sp.* | *Parahaliea sp.* | 97.67 | 0 | 3097080 | 132787 | 0.52 | 2753 | JBKEAX000000000 |
| VLC_044 | *Lachnospiraceae sp.* | *Lachnospiraceae bacterium* | 97.56 | 0.4 | 3210331 | 107442 | 0.41 | 2904 | JBKEAY000000000 |
| VLC_045 | GCA-002748415 sp. | *Actinomycetales bacterium* | 97.46 | 2.18 | 5307799 | 96063 | 0.69 | 4577 | JBKEAZ000000000 |
| VLC_046 | *Lachnospirales sp.* | *Lachnospirales bacterium* | 97.4 | 3.71 | 2366842 | 18575 | 0.29 | 2200 | JBKEBA000000000 |
| VLC_047 | *Shewanella algae* | *Shewanella algae* | 97.36 | 2.62 | 4717750 | 17112 | 0.53 | 4536 | JBKEBB000000000 |
| VLC_048 | CADEDH01 sp. | *Acidimicrobiales bacterium* | 97.29 | 4.92 | 4700912 | 65495 | 0.71 | 4273 | JBKEBC000000000 |
| VLC_049 | UBA2834 sp. | *Candidatus Saccharimonadales bacterium* | 97.23 | 0.92 | 948797 | 185233 | 0.47 | 976 | JBKEBD000000000 |
| VLC_050 | UPXZ01 sp. | *Paludibacteraceae bacterium* | 97.2 | 0.2 | 2813850 | 43019 | 0.39 | 2273 | JBKEBE000000000 |
| VLC_051 | *Bacteroidales sp.* | *Bacteroidales bacterium* | 97.16 | 2.81 | 3199891 | 34842 | 0.4 | 2651 | JBKEBF000000000 |
| VLC_052 | *Pikeienuella sp.* | *Pikeienuella sp.* | 97.15 | 2.5 | 3586985 | 10428 | 0.67 | 3694 | JBKEBG000000000 |
| VLC_053 | *Lachnospirales sp.* | *Lachnospirales bacterium* | 97.08 | 2.09 | 3047668 | 109375 | 0.32 | 2801 | JBKEBH000000000 |
| VLC_054 | *Dysgonomonas sp.* | *Dysgonomonas sp.* | 97.04 | 3.16 | 3730927 | 23472 | 0.38 | 3280 | JBKEBI000000000 |
| VLC_055 | *Mangrovibacterium sp.* | *Mangrovibacterium sp.* | 97.02 | 2.94 | 2890383 | 21225 | 0.45 | 2546 | JBKEBJ000000000 |
| VLC_056 | *Campylobacteraceae sp.* | *Campylobacteraceae bacterium* | 96.93 | 1 | 2030266 | 19232 | 0.33 | 2190 | JBKEBK000000000 |
| VLC_057 | *Lactovum sp.* | *Lactovum sp.* | 96.9 | 0.11 | 1560817 | 25060 | 0.33 | 1571 | JBKEBL000000000 |
| VLC_058 | *Planctomycetaceae sp.* | *Planctomycetaceae bacterium* | 96.58 | 2.85 | 6142505 | 19309 | 0.61 | 5360 | JBKEBM000000000 |
| VLC_059 | *Desulfopila sp.* | *Desulfopila sp.* | 96.58 | 2.05 | 4746059 | 50971 | 0.57 | 4014 | JBKEBN000000000 |
| VLC_060 | *Pseudomonas_A sp.* | *Pseudomonas sp.* | 96.53 | 2.31 | 4122444 | 53131 | 0.61 | 3852 | JBKEBO000000000 |
| VLC_061 | UBA5962 sp. | *Lachnospirales bacterium* | 96.51 | 0.16 | 2481100 | 75763 | 0.31 | 2311 | JBKEBP000000000 |
| VLC_062 | *Vibrio sp.* | *Vibrio sp.* | 96.4 | 1.29 | 4115181 | 41067 | 0.43 | 3884 | JBKEBQ000000000 |
| VLC_063 | *Brevilactibacter sp.* | *Brevilactibacter sp.* | 96.25 | 2.53 | 2854989 | 20774 | 0.68 | 2856 | JBKEBR000000000 |
| VLC_064 | *Brevilactibacter sp.* | *Brevilactibacter sp.* | 96.1 | 4.95 | 3198868 | 28859 | 0.65 | 3118 | JBKEBS000000000 |
| VLC_065 | QQUY01 sp. | *Pyrinomonadaceae bacterium* | 95.94 | 1.69 | 3546989 | 35292 | 0.43 | 3252 | JBKEBT000000000 |
| VLC_066 | *Ruminococcaceae sp.* | *Ruminococcaceae bacterium* | 95.83 | 0.85 | 3663147 | 37667 | 0.57 | 3331 | JBKEBU000000000 |
| VLC_067 | JAAYCI01 sp. | *Ruminococcaceae bacterium* | 95.78 | 5.24 | 3222245 | 8535 | 0.54 | 3222 | JBKEBV000000000 |
| VLC_068 | *Breznakia sp.* | *Breznakia sp.* | 95.7 | 0.27 | 1819740 | 67813 | 0.35 | 1829 | JBKEBW000000000 |
| VLC_069 | *Suipraeoptans sp.* | *Suipraeoptans sp.* | 95.62 | 1.47 | 3642326 | 31472 | 0.37 | 3495 | JBKEBX000000000 |
| VLC_070 | *Alphaproteobacteria sp.* | *Alphaproteobacteria bacterium* | 95.61 | 0.06 | 1997652 | 43768 | 0.39 | 1852 | JBKEBY000000000 |
| VLC_071 | *Leucobacter sp.* | *Leucobacter sp.* | 95.26 | 6.4 | 3809236 | 33535 | 0.67 | 3484 | JBKEBZ000000000 |
| VLC_072 | *Xiphinematobacteraceae sp.* | *Chthoniobacterales bacterium* | 95.23 | 0.96 | 2449084 | 14163 | 0.51 | 2477 | JBKECA000000000 |
| VLC_073 | *Paracoccus sp.* | *Paracoccus sp.* | 95.17 | 5.12 | 4882789 | 23949 | 0.64 | 4906 | JBKECB000000000 |
| VLC_074 | *Anaerorhabdus sp.* | *Anaerorhabdus sp.* | 95.14 | 0.05 | 1509366 | 159192 | 0.3 | 1428 | JBKECC000000000 |
| VLC_075 | *Coprobacillaceae sp.* | *Coprobacillaceae bacterium* | 95.1 | 8.39 | 3533362 | 10294 | 0.3 | 3827 | JBKECD000000000 |
| VLC_076 | UBA660 sp. | *Bacilli bacterium* | 94.94 | 1.57 | 1416169 | 28363 | 0.29 | 1500 | JBKECE000000000 |
| VLC_077 | *Paracoccus sp.* | *Paracoccus sp.* | 94.94 | 3.16 | 3445273 | 18158 | 0.62 | 3706 | JBKECF000000000 |
| VLC_078 | *Arachnia sp.* | *Arachnia sp.* | 94.76 | 1.94 | 2856895 | 30318 | 0.67 | 2801 | JBKECG000000000 |
| VLC_079 | *Oleidesulfovibrio sp.* | *Oleidesulfovibrio sp.* | 94.58 | 0.05 | 3836873 | 129041 | 0.52 | 3446 | JBKECH000000000 |
| VLC_080 | UPXZ01 sp. | *Paludibacteraceae bacterium* | 94.56 | 0.8 | 2906115 | 30104 | 0.41 | 2635 | JBKECI000000000 |
| VLC_081 | *Mangrovibacterium sp.* | *Mangrovibacterium sp.* | 94.47 | 0.82 | 2416906 | 91719 | 0.41 | 1997 | JBKECJ000000000 |
| VLC_082 | *Beutenbergiaceae sp.* | *Beutenbergiaceae bacterium* | 94.39 | 3.62 | 2584929 | 41770 | 0.64 | 2479 | JBKECK000000000 |
| VLC_083 | UBA7332 sp. | *Bacteroidales bacterium* | 94.37 | 0.26 | 2985209 | 10612 | 0.43 | 2787 | JBKECL000000000 |
| VLC_084 | *Dysgonomonas sp.* | *Dysgonomonas sp.* | 94.25 | 2.51 | 4055402 | 67715 | 0.37 | 3591 | JBKECM000000000 |
| VLC_085 | *Bacteroides graminisolvens* | *Bacteroides graminisolvens* | 94.25 | 0.29 | 3422346 | 17286 | 0.41 | 3031 | JBKECN000000000 |
| VLC_086 | *Microbacterium sp.* | *Microbacterium sp.* | 94.24 | 2.95 | 3283295 | 20919 | 0.69 | 3306 | JBKECO000000000 |
| VLC_087 | JAEUMD01 sp. | *Hyphomicrobiaceae bacterium* | 94.14 | 1.77 | 3389401 | 7188 | 0.57 | 3469 | JBKECP000000000 |
| VLC_088 | *Dysgonomonas sp.* | *Dysgonomonas sp.* | 94.06 | 9.85 | 4790442 | 26753 | 0.37 | 4464 | JBKECQ000000000 |
| VLC_089 | *Azonexus sp.* 900549295 | *Azonexus sp.* | 93.98 | 0.47 | 2421938 | 11041 | 0.64 | 2523 | JBKECR000000000 |
| VLC_090 | *Dysgonomonas sp.* 002482385 | *Dysgonomonas sp.* | 93.94 | 4.67 | 4337223 | 42762 | 0.4 | 3633 | JBKECS000000000 |
| VLC_091 | *Bacteroides thetaiotaomicron* | *Bacteroides thetaiotaomicron* | 93.91 | 1.48 | 6190603 | 157842 | 0.43 | 4870 | JBKECT000000000 |
| VLC_092 | *Anaerotignum sp.* | *Anaerotignum sp.* | 93.77 | 2 | 1868791 | 6874 | 0.38 | 2004 | JBKECU000000000 |
| VLC_093 | *Alphaproteobacteria sp.* | *Alphaproteobacteria bacterium* | 93.73 | 2.23 | 2123149 | 52450 | 0.4 | 2010 | JBKECV000000000 |
| VLC_094 | *Tropicimonas sp.* | *Tropicimonas sp.* | 93.64 | 8.53 | 4573370 | 35008 | 0.66 | 4433 | JBKECW000000000 |
| VLC_095 | *Bacteroides xylanisolvens* | *Bacteroides xylanisolvens* | 93.62 | 9 | 7346975 | 57526 | 0.41 | 5841 | JBKECX000000000 |
| VLC_096 | *Anaerocolumna sp.* | *Anaerocolumna sp.* | 93.4 | 0.03 | 3904114 | 34864 | 0.34 | 3459 | JBKECY000000000 |
| VLC_097 | *Mangrovibacterium sp.* | *Mangrovibacterium sp.* | 93.34 | 4.53 | 4239118 | 21822 | 0.46 | 3755 | JBKECZ000000000 |
| VLC_098 | JAAYCI01 *sp.* | *Ruminococcaceae bacterium* | 93.27 | 1.62 | 2977434 | 34362 | 0.47 | 2898 | JBKEDA000000000 |
| VLC_099 | *Parachlamydiaceae sp.* | *Parachlamydiaceae bacterium* | 93.21 | 2.44 |  | 13788 | 0.46 | 2549 | JBKEDB000000000 |
| VLC_100 | *Halodesulfovibrio sp.* | *Halodesulfovibrio sp.* | 92.99 | 2.52 | 4128835 | 21213 | 0.57 | 3839 | JBKEDC000000000 |
| VLC_101 | *Bacteroidales sp.* | *Bacteroidales bacterium* | 92.9 | 0.71 | 2610130 | 31609 | 0.41 | 2378 | JBKEDD000000000 |
| VLC_102 | *Microtrichaceae sp.* | *Microtrichaceae bacterium* | 92.75 | 1.76 | 3654096 | 19389 | 0.67 | 3534 | JBKEDE000000000 |
| VLC_103 | *Desertimonas sp.* | *Desertimonas sp.* | 92.52 | 2.37 | 3977315 | 20184 | 0.7 | 4014 | JBKEDF000000000 |
| VLC_104 | *Moheibacter sp.* | *Moheibacter sp.* | 92.33 | 4.06 | 2308769 | 20990 | 0.34 | 2334 | JBKEDG000000000 |
| VLC_105 | *Microbacterium sp.* | *Microbacterium sp.* | 92.16 | 3.43 | 2084114 | 29515 | 0.69 | 2094 | JBKEDH000000000 |
| VLC_106 | *Tropicimonas sp.* | *Tropicimonas sp.* | 92.1 | 1.36 | 3493884 | 28904 | 0.65 | 3404 | JBKEDI000000000 |
| VLC_107 | *Xiphinematobacteraceae sp.* | *Chthoniobacterales bacterium* | 91.92 | 0.01 | 2466470 | 13064 | 0.49 | 2476 | JBKEDJ000000000 |
| VLC_108 | *Mangrovibacterium sp.* | *Mangrovibacterium sp.* | 91.89 | 0.91 | 3103440 | 26005 | 0.39 | 2508 | JBKEDK000000000 |
| VLC_109 | *Paracoccus sp.* | *Paracoccus sp.* | 91.74 | 1.52 | 2882721 | 19942 | 0.56 | 3046 | JBKEDL000000000 |
| VLC_110 | *Castellaniella sp.* | *Castellaniella sp.* | 91.71 | 1.23 | 2831004 | 111946 | 0.65 | 2619 | JBKEDM000000000 |
| VLC_111 | *Demequina sp.* | *Demequina sp.* | 91.64 | 9.93 | 2973876 | 7059 | 0.69 | 3208 | JBKEDN000000000 |
| VLC_112 | Ga0077555 *sp.* | *Hyphomicrobiaceae bacterium* | 91.48 | 5.57 | 4936968 | 7267 | 0.63 | 5072 | JBKEDO000000000 |
| VLC_113 | *Paracoccus sp.* | *Paracoccus sp.* | 91.47 | 9.94 | 4315115 | 7498 | 0.66 | 4838 | JBKEDP000000000 |
| VLC_114 | *Rhodoblastus sp.* | *Rhodoblastus sp.* | 91.27 | 0.51 | 3214082 | 25634 | 0.64 | 3098 | JBKEDQ000000000 |
| VLC_115 | UBA3375 sp. | *Mycoplasmatales bacterium* | 91.22 | 7.89 | 1468599 | 19897 | 0.26 | 1484 | JBKEDR000000000 |
| VLC_116 | *Tenacibaculum sp.* | *Tenacibaculum sp.* | 91.01 | 0 | 2573578 | 38858 | 0.33 | 2136 | JBKEDS000000000 |
| VLC_117 | UBA3375 *sp.* | *Mycoplasmatales bacterium* | 90.92 | 7.77 | 991124 | 27701 | 0.27 | 1069 | JBKEDT000000000 |
| VLC_118 | UJ101 *sp.* | *Flavobacteriales bacterium* | 90.89 | 0.12 | 1629569 | 14643 | 0.31 | 1568 | JBKEDU000000000 |
| VLC_119 | *Qingshengfaniella sp.* | *Qingshengfaniella sp.* | 90.77 | 0.95 | 3204093 | 34211 | 0.64 | 3071 | JBKEDV000000000 |
| VLC_120 | *Parahaliea sp.* | *Parahaliea sp.* | 90.62 | 0.49 | 3534554 | 5230 | 0.63 | 3733 | JBKEDW000000000 |
| VLC_121 | UBA5026 *sp.* | *Bacilli bacterium* | 90.55 | 0.09 | 977605 | 105097 | 0.25 | 1051 | JBKEDX000000000 |
| VLC_122 | *Beutenbergiaceae sp.* | *Beutenbergiaceae bacterium* | 90.46 | 2.87 | 2989408 | 71829 | 0.66 | 2855 | JBKEDY000000000 |
| VLC_123 | CADEDH01 *sp.* | *Acidimicrobiales bacterium* | 90.4 | 2.65 | 4276914 | 22849 | 0.71 | 4094 | JBKEDZ000000000 |
| VLC_124 | *Lacrimispora sphenoides_A* | *Lacrimispora sphenoides* | 90.32 | 4.45 | 5235146 | 14751 | 0.44 | 5166 | JBKEEA000000000 |
| VLC_125 | *Erysipelotrichaceae sp.* | *Erysipelotrichaceae bacterium* | 90.3 | 2.23 | 1345357 | 24573 | 0.31 | 1367 | JBKEEB000000000 |
| VLC_126 | *Draconibacterium sp.* | *Draconibacterium sp.* | 90.13 | 6.44 | 5014037 | 6542 | 0.43 | 4916 | JBKEEC000000000 |
| VLC_127 | *Macellibacteroides fermentans* | *Macellibacteroides fermentans* | 90.09 | 0.33 | 4127497 | 51099 | 0.41 | 3425 | JBKEED000000000 |
| VLC_128 | UBA660 sp. | *Bacilli bacterium* | 90.08 | 1.29 | 872413 | 40567 | 0.24 | 971 | JBKEEE000000000 |
| VLC_129 | *Kluyvera sp.* | *Kluyvera sp.* | 90.07 | 2.86 | 5138619 | 74372 | 0.56 | 4742 | JBKEEF000000000 |

### Beta diversity (community structure) is correlated with crab host genus and terrestrial grade

For bacterial communities (K2/B and MAG), host genus was significantly correlated with beta diversity (PERMANOVA, p < 0.05), while terrestrial grade was significantly correlated with MAG beta diversity (PERMANOVA, p < 0.05), but marginally non significant for K2/B beta diversity (p = 0.055). In both models, host genus explained more variation (K2/B: 12.95%; MAG: 19.64%) than grade (K2/B: 6.13%; MAG: 6.03%) (Figure 3, Figure S1, Table S2). *Post hoc* pairwise PERMANOVA contrasts for bacterial communities showed that host genus differences (for both K2/B and MAG datasets) were driven by significant differences between a few host genera, specifically *Petrolisthes* vs. *Birgus*, *Tuerkayana* and *Gecarcoidea* (p < 0.05), while *post hoc* pairwise PERMANOVA contrasts were significantly different across all grades for MAGs except I vs. II and all grades except III vs. IV for K2/B bacteria.

**Figure 3.**
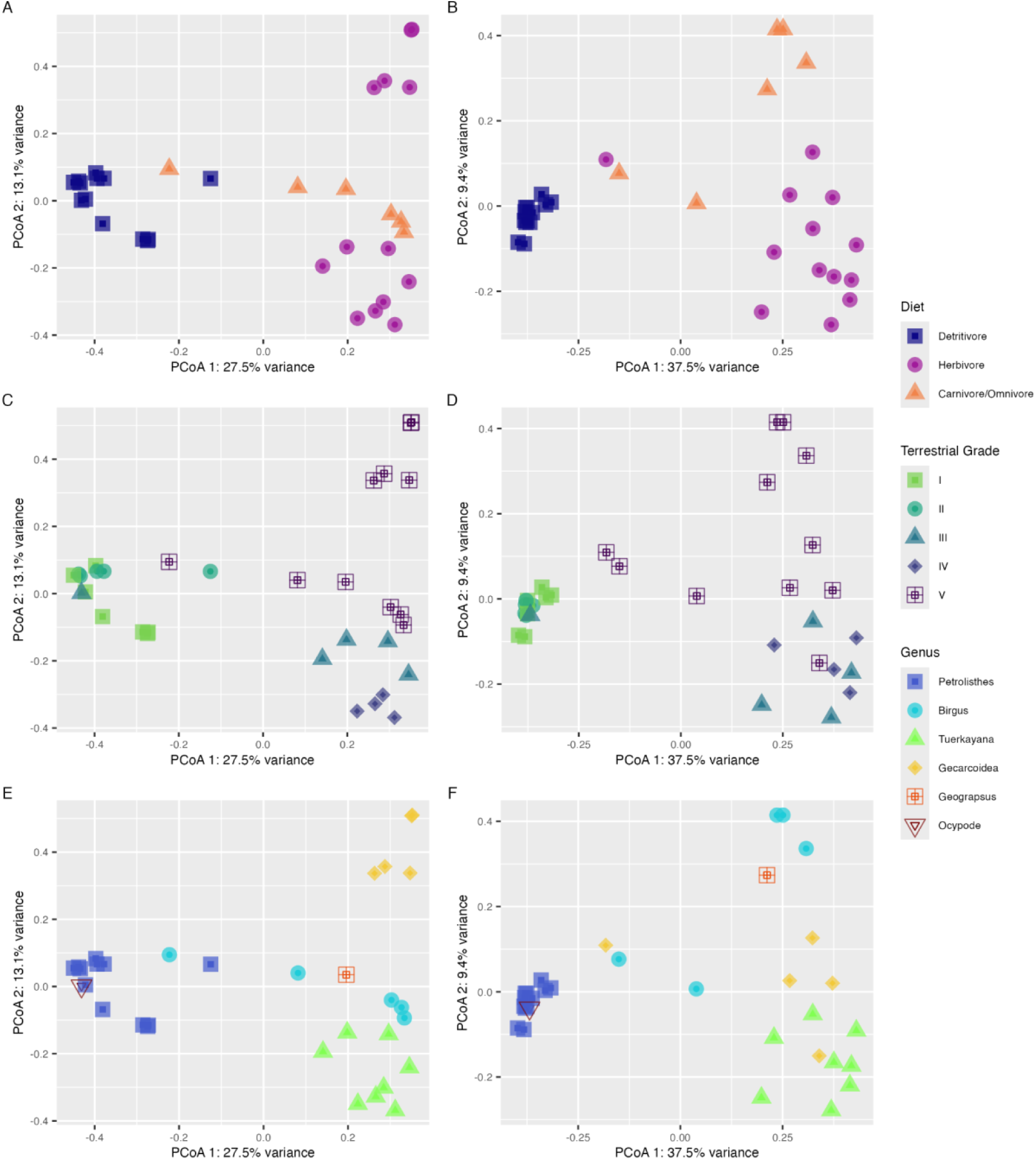
Bacterial beta diversity across diet, terrestrial grades and crab genera. Principal-coordinate analysis (PCoA) visualization of Bray Curtis dissimilarities of coverM counts of MAG abundance showing individual samples designated as points with colors and shapes representing (A) diet (detritivore, herbivore, carnivore/omnivore), (C) terrestrial grade (I-V), and (E) crab genera. Principal-coordinate analysis (PCoA) visualization of Bray Curtis dissimilarities of Kraken2/Bracken counts of bacterial abundance showing individual samples designated as points with colors and shapes representing (B) diet, (D) terrestrial grade, and (F) crab genera.

We also tested for dispersion (variance) differences between these groups. We found that MAGs had significant differences in dispersion for both host genus and grade (betadisper, p < 0.05, Table S2), with *post hoc* Dunn tests revealing that these differences were driven by differences between *Tuerkayana* vs. *Geograpsus*, and *Ocypode*, and differences between grade V vs. grade I and grade II (Table S3). We found that while the overall test indicated significant differences in dispersion between terrestrial grades for the K2/B bacteria, *post hoc* Dunn tests identified no pairwise contrasts that were significantly different (p > 0.05).

For fungal communities (K2/B), we found that host genus and grade were significantly correlated with beta diversity (PERMANOVA, p < 0.05) (Figure S1). *Post hoc* pairwise contrasts revealed that this was driven by differences between *Petrolisthes* vs. *Birgus* & *Tuerkayana* and *Tuerkayana* vs. *Birgus,* as well as grade II with most other grades. We also found significant differences in dispersion across host genera (p < 0.05), but not across grades (p > 0.05), with *post hoc* Dunn tests showing that these were driven by *Geograpsus* and *Ocypode* vs. *Petrolisthes*, *Tuerkayana*, and *Birgus*.

For archaeal communities (K2/B) (Figure S1), there were no significant differences in beta diversity (PERMANOVA, p > 0.05) and also no significant differences in dispersion (betadisper, p > 0.05).

### Relative abundance analyses identified potential key microbial taxa that differ across crab genera, diet, and terrestrial grade

In terms of microbial community composition based on K2/B - for bacteria, the most prevalent phyla on average were Pseudomonadota (i.e., Proteobacteria; 54.40%), Bacillota (i.e., Firmicutes; 15.70%), Bacteroidota (i.e., Bacteroidetes; 9.86%), Actinomycetota (i.e., Actinobacteria; 9.53%), and Mycoplasmatota (i.e., Tenericutes; 2.24%). While at the genus level we observed the dominant genera to be *Pseudomonas* (Class Gammaproteobacteria), *Vibrio* (Class Gammaproteobacteria), *Citrobacter* (Class Gammaproteobacteria), *Escherichia* (Class Gammaproteobacteria), and uncultured bacteria (Figure S2). Among Archaea, the dominant phyla were Euryarchaeota (59.50%) and unclassified archaea (27.50%), while the dominant genera were *Methanosarcina* and uncultured archaea (Figure S3). For fungi, the most abundant phyla were Ascomycota (74.50%), Basidiomycota (14.40%), and Mucoromycota (6.16%), while *Saccharomyces*, *Aspergillus*, *Fusarium*, *Rhizophagus*, and uncultured fungi were prominent genera (Figure S4). We aimed to explore variations in the relative abundance of abundant microbial genera across different diets, terrestrial grades, and host genera, with a focus on cross-referencing with known lignin and cellulose degraders.

Intriguingly, certain bacterial genera displayed significant differences in relative abundance across multiple categories (Figure S5, Table S4-S5). Notably, *Citrobacter* (Gammaproteobacteria), *Escherichia* (Class Gammaproteobacteria), *Paenibacillus* (Bacilli), and *Paracoccus* (Alphaproteobacteria) demonstrated variations across all three categories (diet, terrestrial grade, and host genera). *Citrobacter* and *Paracoccus* were more relatively abundant in host genera at higher grades of terrestriality, while *Paenibacillus* and *Escherichia* were more relatively abundant in host genera at lower grades of terrestriality. No archaea were differentially relatively abundant across all three groups in union, but across diet, we found that *Methanobrevibacter* was more relatively abundant in carnivores, while *Natrinema* was more relatively abundant in both detritivores and herbivores (with slightly higher relative abundance in detritivores). Across host genera, we also observed significant differences in unclassified *Candidatus* Methanoperedenaceae relative abundance in *Birgus*. For fungi, *Trametes* relative abundance was significantly different across all three categories and was also more relatively abundant in host genera at higher terrestrial grades.

We also used CoverM to explore whether relative abundance of bacterial MAGs would recapitulate the trends discussed above or expose new patterns in relative abundance across different diets, terrestrial grades, and host genera. The most prevalent MAG phyla based on average relative abundance were Bacteroidota (i.e., Bacteroidetes; 44.80%), Pseudomonadota (i.e., Proteobacteria; 34.50%), Bacillota (i.e., Firmicutes; 11.56%), and Actinomycetota (i.e., Class Actinobacteria; 5.56%). Importantly, we could only assemble MAGs for one *Citrobacter* and four *Paracoccus* species alongside fifteen Bacilli MAGs (although *Paenibacillus* was notably absent), and eight other Enterobacterales (Class Gammaproteobacteria) MAGs were present but *Escherichia* was not, somewhat limiting our comparisons between approaches. There are several possible reasons for discrepancies between the MAG and the read-based analyses, including different naming conventions between databases [63, 64], and the overall limitations of read-based taxonomic classifiers [65], which can be complicated by low coverage or host-dominated metagenomic sequencing. But generally, we found that several MAG genera had significant differences in relative abundance across all three categories. These included *Citrobacter* and *Paracoccus*, which had been detected in the K2/B approach, as well as *Kluyvera* (Class Gammaproteobacteria) and *Anaerorhabdus* (Bacilli), which both belong to groups highlighted in the K2/B approach. Furthermore, there were significant differences in new groups not identified in the K2/B approach, including members of novel Alphaproteobacteria bacterium, *Demequina* (Actinomycetia), *Marinicella* (Class Gammaproteobacteria), *Moheibacter* (Bacteroidota), and UJ101 (Bacteroidota).

Specifically, novel Alphaproteobacteria bacteria were significantly less relatively abundant in detritivores but were more relatively abundant in land crabs at higher terrestrial grades. We also saw this pattern for *Anaerorhabdus* (Bacilli), which stands in contrast to our finding for *Paenibacillus* (Bacilli) using the K2/B approach. Consistent with our K2/B results, we also found that *Citrobacter* (Class Gammaproteobacteria) and *Paracoccus* (Alphaproteobacteria) were significantly more relatively abundant in herbivores and in land crabs at higher terrestrial grades, and although they were not detected using the K2/B approach, *Moheibacter* (Bacteroidota) and *Demequina* (Actinomycetia) also reflected this pattern. Finally, *Kluyvera* (Enterobacterales) and UJ101 (Bacteriodota) were both more relatively abundant in detritivores and in crabs from lower terrestrial grades (matching the pattern we found for *Escherichia* (Enterobacterales: Class Gammaproteobacteria) using the K2/B approach), while *Marinicella* (Class Gammaproteobacteria) was only more relatively abundant in crabs from lower grades.

### Predicted metabolic functional differences in the gut microbiome between crabs from different terrestrial grades

Broad-scale functional analysis of metagenomic samples, which are predictions of functions based on sequence similarity, revealed that samples from crab genera from lower terrestrial grades (I-II) lack complete complex carbon degradation pathways, including almost a complete absence of hemicellulose de-branching (i.e. alpha-L-arabinofuranosidase [EC:3.2.1.55], beta-galactosidase [EC:3.2.1.23], and alpha-L-rhamnosidase [EC:3.2.1.40]) and endohemicellulase (i.e. endo-1,4-beta-xylanase [EC:3.2.1.8] and xylose isomerase [EC:5.3.1.5]) proteins, both of which are present in most genera from higher terrestrial grades (IV-V). Cellulose degrading enzymes (i.e., endo-beta-1,4-glucanase [EC:3.2.1.4], cellulose 1,4-beta-cellobiosidase [EC:3.2.1.91], and beta-glucosidase [EC:3.2.1.21]) were also poorly represented in metagenomic samples collected from crabs from grades I-Il (Figure 4). For grade III, *Ocypode* samples tended to reflect patterns seen in crabs from lower grades, while *Tuerkayana celeste* samples largely matched patterns observed in the more terrestrially adapted crab species, implying that specific diet type may play a critical role in determining gut microbial assembly and function within grades.

**Figure 4.**
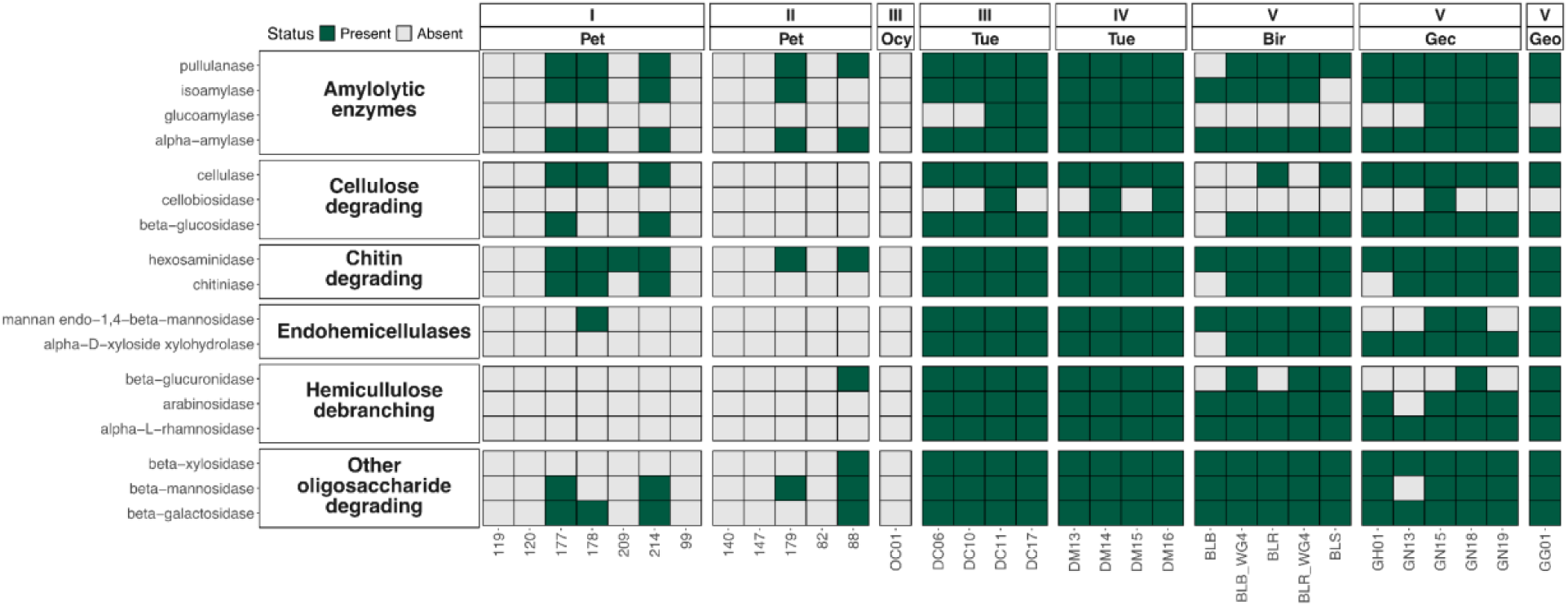
Complex carbon degradation pathway detection. METABOLIC (Zhou et al. 2022) was used to profile the detection of genes involved in complex carbon degradation across assembled bacterial contigs from metagenomic samples. The heatmap summarizes 17 horizontal entries based on key functional gene families covering 6 function categories. Cells display the detection (presence: green/absence: gray) of these genes across samples which are grouped by terrestrial grade (I-V) and crab genera. Crab genera are noted using a three-letter code (Pet = *Petrolisthes*, Ocy = *Ocypode*, Tue = *Tuerkayana*, Bir = *Birgus*, Gec = *Gecarcoidea*, Geo = *Geograpsus*) for visualization purposes. Detailed KEGG KO identifier IDs, gene names, and protein information for each function category used to generate the heatmap are described in Supplementary Table S7.

We used Random Forest classification to predict crab diet based on complex carbon degradation pathway gene counts. On the 30% hold-out test set (n = 9), the model achieved an overall accuracy of 85.71% on the test set, with a Kappa value of 0.78, reflecting substantial agreement between predicted and observed diet categories. Consistent with sample representation across diet type, the confusion matrix showed that the model did well predicting detritivores (n =13) and herbivores (n = 13), which were well-sampled, but was less reliable for the undrerrepresented carnivore/omnivore categories (n = 6 combined) (Table S4). Amylolytic enzymes were the most important predictor based on feature importance for diet classification, followed by chitin-degrading enzymes and hemicellulose de-branching enzymes. (Table S6).

We detected 266 unique CAZyme families across all samples, 12,988 gene-family assignments: 53.0% of detected CAZyme domains belonged to glycoside hydrolases (GHs), 27.6% to glycosyltransferases (GTs), 7.58% to carbohydrate esterases (CEs), 5.18% to carbohydrate-binding modules (CBMs), 3.56% to polysaccharide lyases (PLs), and 3.14% to auxiliary activities (AAs). Restricting to families linked to lignocellulose degradation and only considering complete genes yielded 3,306 gene-family hits across 34 families, with at least one hit in all 129 MAGs.

Among lignocellulose-related families, GH2 was the most abundant (536 genes), while GH3 and GH13 were the most widely distributed, detected in 84% and 80% of MAGs respectively (Figure S6). GH13 was detected in most MAGs of every major class. Most Bacteroidia (specifically Bacteriodales) MAGs were also enriched for cellulases (GH5 and GH30), hemicellulases (GH29, CE1, and CE4), beta glucosidases (GH2 and GH3), beta-1-3-glucanase (GH16), and glycosyl hydrolase (GH20) (Figure 5). Clostridia MAGs were enriched for cellulases (GH5 and GH30), beta glucosidases (GH2 and GH3), and hemicellulase (CE4) (Figure S7). The majority of Bacteroidia and Clostridia MAGs with CAZyme enrichment were collected from *Tuerkayana* samples, and interestingly, the enriched CAZyme repertoire in both bacterial classes was highly concordant.

**Figure 5.**
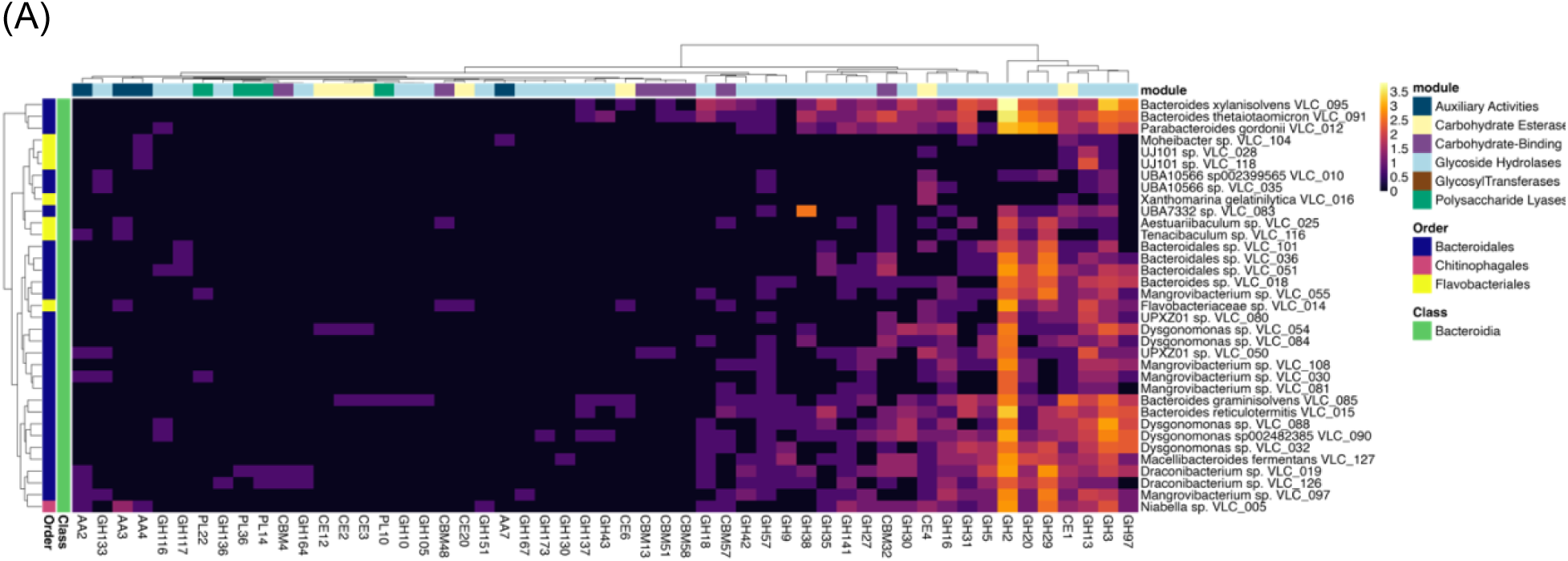
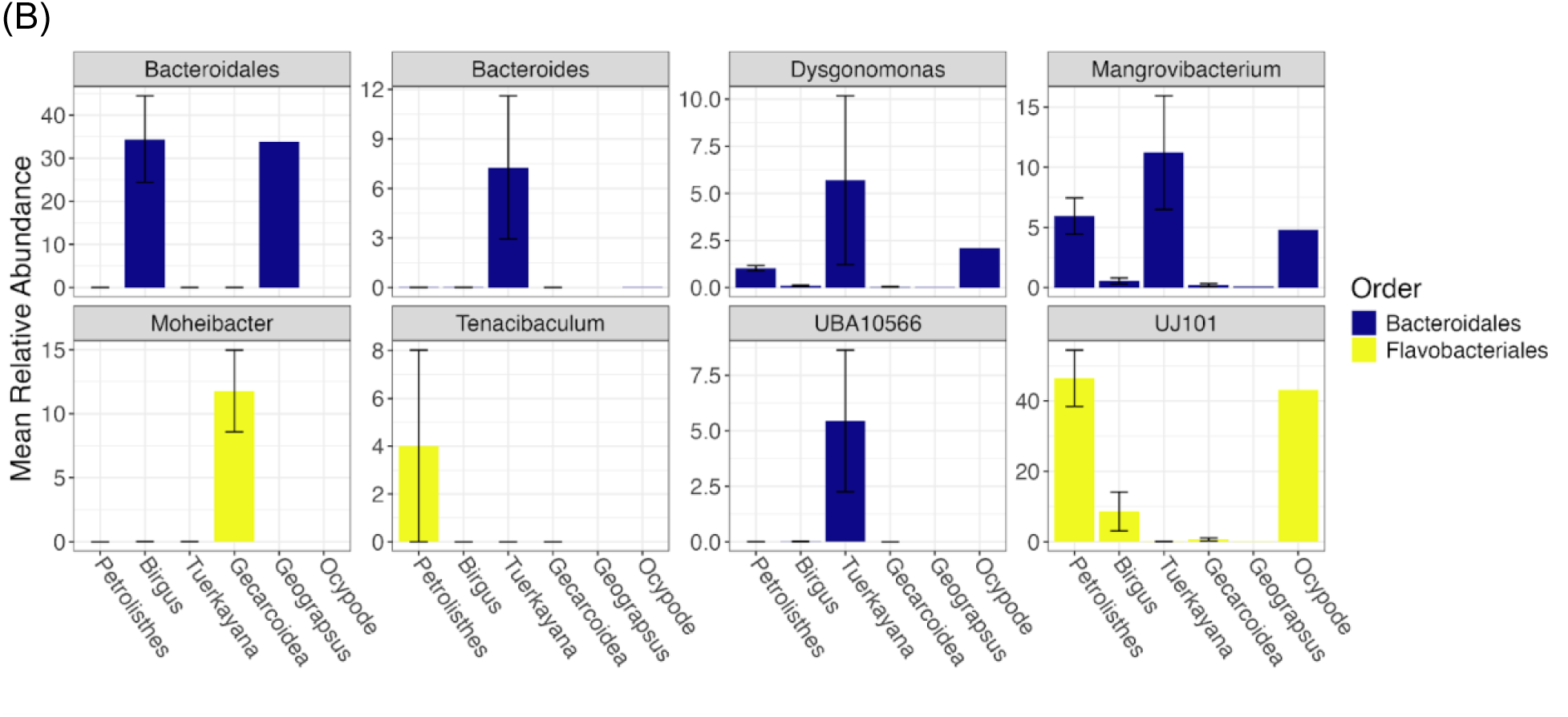
Predicted CAZyme domain abundance across Bacteroidia. (A) DRAM was used to annotate CAZyme domains in the proteome of MAGs. A heatmap displaying the log-transformed counts of CAZyme domains with potential lignocellulose-related functions for members of the Bacteroidia. Specific CAZyme modules are denoted across the top, while MAG taxonomic order and class information are denoted on the left. Further, the dendrogram to the left of the heatmap clusters the MAGs by similarity in counts between the different CAZyme domains, while the dendrogram above clusters CAZymes based on similarity in counts between the different MAGs. (B) Relative abundance of MAGs in the Bacteroidia across host crab genera with a mean relative abundance >1% are displayed as a barplot. Bars are split by taxonomic genus, colored by predicted order and error bars represent standard error. Clades are shown to the lowest assignable taxonomic classification.

Actinomycetia MAGs, which were most abundant in *Gecarcoidea* samples, were enriched for cellobiose dehydrogenases (AA3), laccase (AA1), beta-1-3-glucanase (GH16), and beta glucosidase (GH3) (Figure S8). Alphaproteobacteria MAGs, which were most relatively abundant in *Gecarcoidea* (especially *Paracoccus*) or *Tuerkayana* samples, were highly enriched for cellobiose dehydrogenases (AA3), along with laccase (AA1), vanillyl alcohol oxidase (AA4), and hemicellulase (CE4) enrichment to a lesser extent (Figure 6).

**Figure 6.**
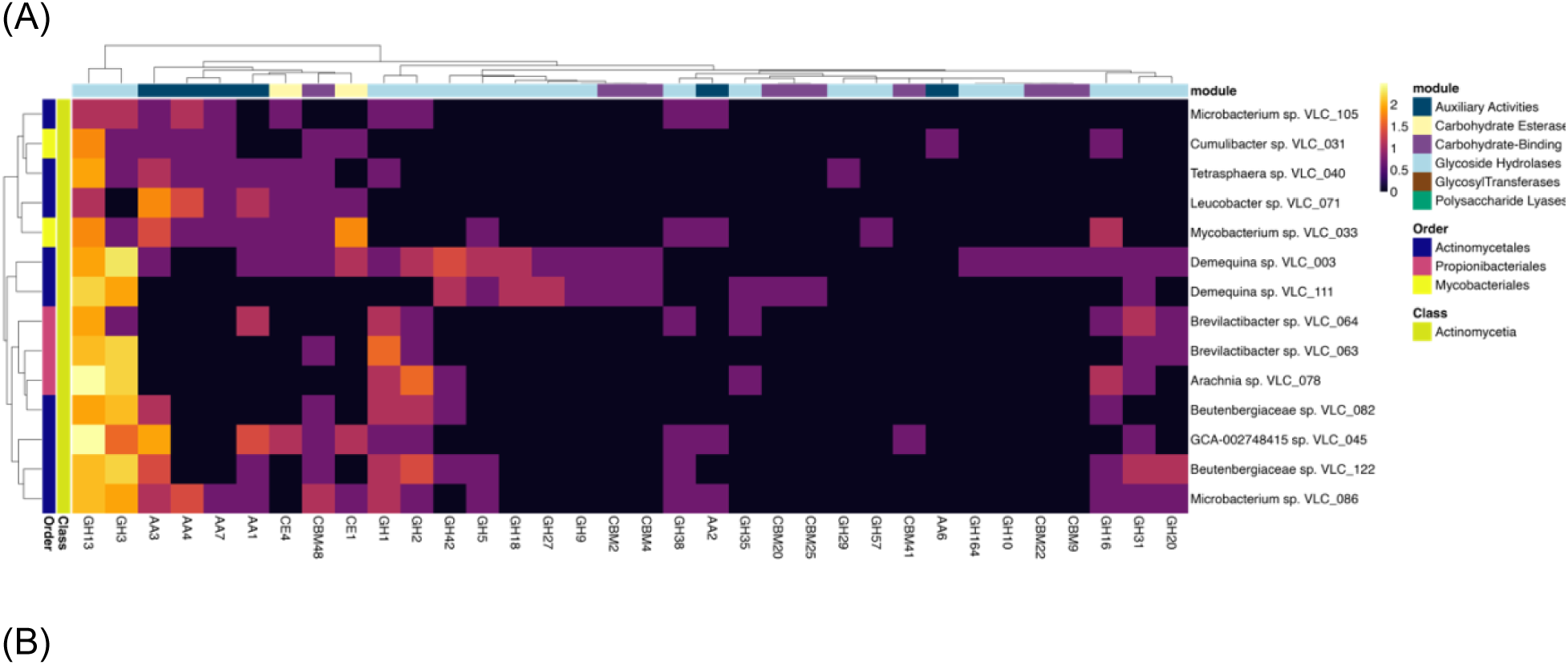
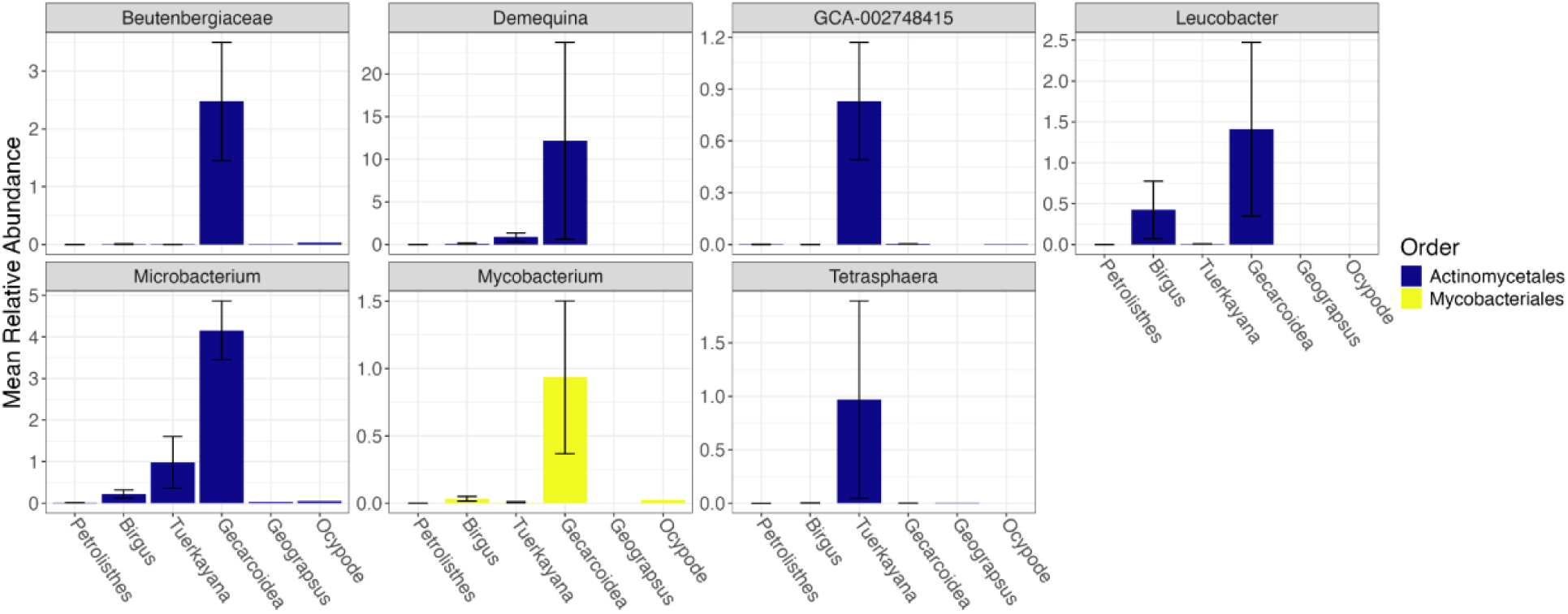
Predicted CAZyme domain abundance across Actinomycetia. (A) DRAM was used to annotate CAZyme domains in the proteome of MAGs. A heatmap displaying the log-transformed counts of CAZyme domains with potential lignocellulose-related functions for members of the Actinomycetia. Specific CAZyme modules are denoted across the top, while MAG taxonomic order and class information are denoted on the left. Further, the dendrogram to the left of the heatmap clusters the MAGs by similarity in counts between the different CAZyme domains, while the dendrogram above clusters CAZymes based on similarity in counts between the different MAGs. (B) Relative abundance of MAGs in the Actinomycetia across host crab genera with a mean relative abundance >0.1% are displayed as a barplot. Bars are split by taxonomic genus, colored by predicted order and error bars represent standard error. Clades are shown to the lowest assignable taxonomic classification.

Class Gammaproteobacteria MAGs, of which some bacterial genera were abundant in *Birgus* (i.e. *Citrobacter*, *Denitromonas*, *Ottowia*) while others were most abundant in crabs from lower grades (e.g. *Kluyvera* and *Marinicella*), were enriched in cellobiose dehydrogenases (AA3), vanillyl alcohol oxidase (AA4), hemicellulases (CE1, and CE4), laccase (AA1), cellulase (GH5), and beta glucosidase (GH1) (Figure 7). Finally, Bacilli, which were abundant in *Gecarcoidea* or *Geograpsus* samples, were enriched for beta glucosidase (GH1) (Figure S9).

**Figure 7.**
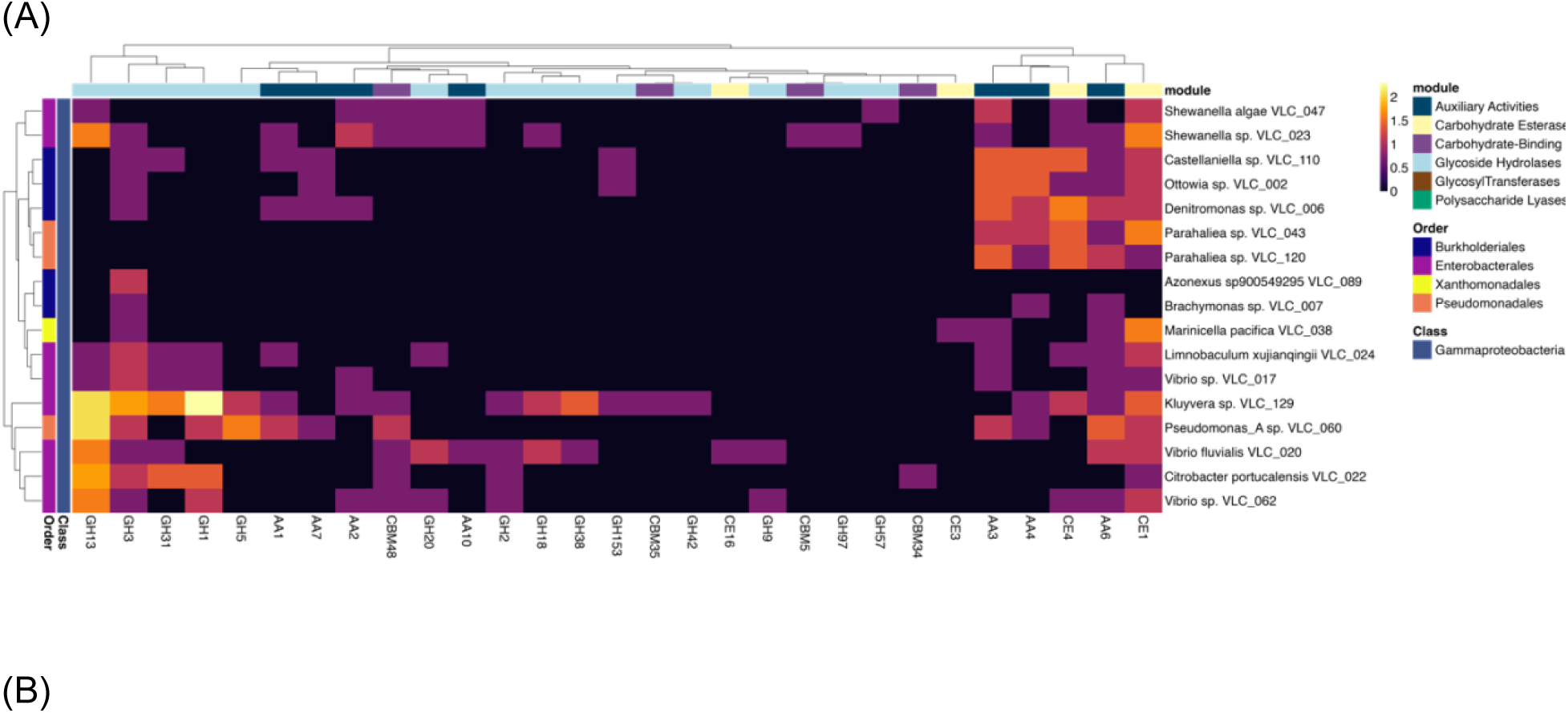
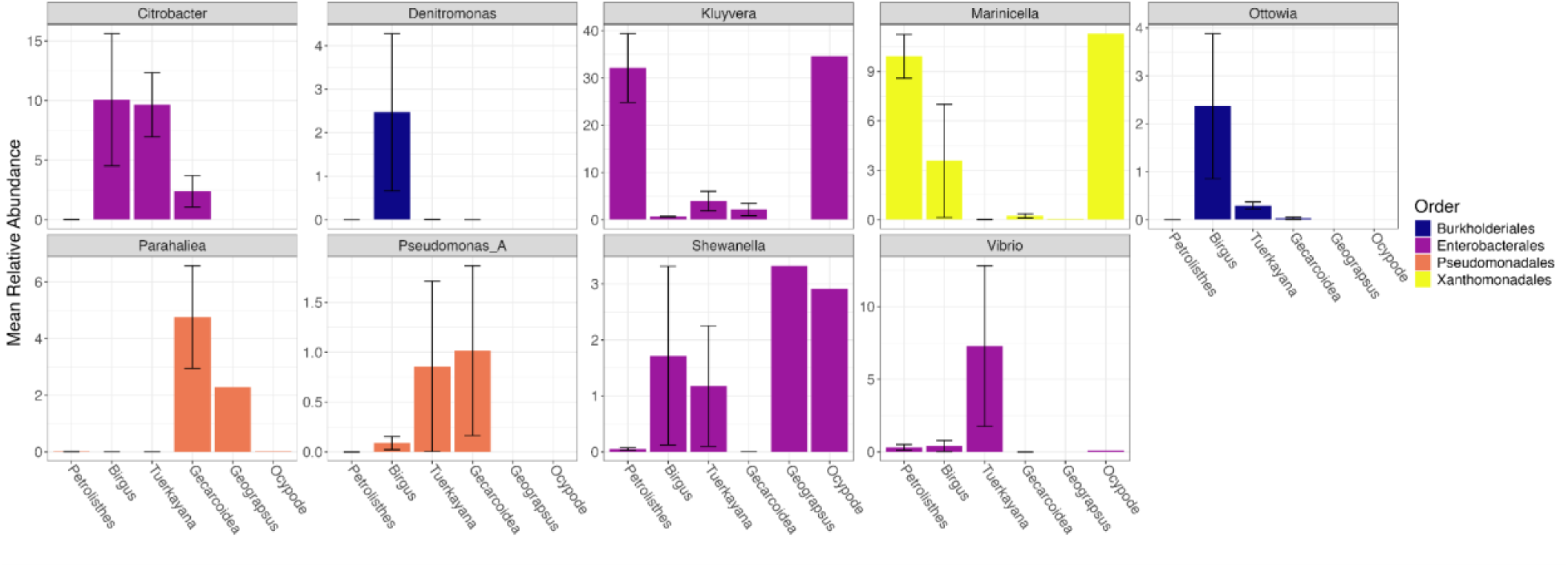
Predicted CAZyme domain abundance across Gammaproteobacteria. (A) DRAM was used to annotate CAZyme domains in the proteome of MAGs. A heatmap displaying the log-transformed counts of CAZyme domains with potential lignocellulose-related functions for members of the class Gammaproteobacteria. Specific CAZyme modules are denoted across the top, while MAG taxonomic order and class information are denoted on the left. Further, the dendrogram to the left of the heatmap clusters the MAGs by similarity in counts between the different CAZyme domains, while the dendrogram above clusters CAZymes based on similarity in counts between the different MAGs. (B) Relative abundance of MAGs in the class Gammaproteobacteria across host crab genera with a mean relative abundance >0.1% are displayed as a barplot. Bars are split by taxonomic genus, colored by predicted order and error bars represent standard error. Clades are shown to the lowest assignable taxonomic classification.

## Discussion

Our comprehensive analysis of the gut microbiome of crabs across a gradient of terrestriality from intertidal regions to forested habitats reveals that host identity (i.e. genus) and degree of terrestriality were both important determinants of microbiome composition. Further, we constructed a foundational genomic resource of 129 bacterial MAGs, including genomes from lignin-degrading bacteria whose relative abundance differed across crab genera and terrestrial grades. We discovered that when looking at specific MAGs, the majority of gut microbiota appeared to have CAZyme domains putatively involved in lignocellulose degradation, regardless of degree of host terrestriality. However, gut microbiomes of crabs from higher terrestrial grades had more complete complex carbon degradation pathways than those from lower grades, indicating an important shift in gut function with habitat. Furthermore, different crab genera had distinct microbial communities in their guts, albeit with similar functional repertoires, suggesting functional convergence in the gut microbiomes of these crabs. Altogether, these results imply that beneficial gut microbes may have convergent roles in lignocellulose degradation, but that these communities are primarily structured based on host identity.

### Gut microbiome composition is largely driven by crab host identity

The variation in microbial community composition was best explained by host genus, followed by terrestrial grade. We infer that this means both phylogenetic relatedness (which includes some variation due to diet) and specific terrestrial habitats and terrestrially adaptive traits (which are both encompassed by the terrestrial grade designations), are also important driving factors of crab gut microbial communities. Previous studies have shown that the microbiome of mangrove-dwelling crabs is distinct from the community of free-living microbes found in their local environment [16], and studies in aquatic invertebrates more generally (e.g., snails, aquatic crabs, etc.) have also observed that gut microbial communities in these clades are distinctly structured and show some specificity by host identity [17, 66, 67].

Additionally, the community-level patterns seen here are similar to what has been recently reported for brachyuran crabs in the family Sesarmidae, where host crab species, followed by local environmental conditions, primarily drive the composition of gut microbial communities [16]. We observed a similar impact of host crab genus (e.g., 12.95% for K2/B and 19.64% for MAGs vs. 15.8-25.3%). Tsang et al. [16] found that terrestriality had only a relatively small effect (1.0-1.6%) on the microbial community composition, whereas we found it to have a larger effect (6.03-6.13%), likely due to our inclusion of species from higher terrestrial grades (i.e., the present work sampled crabs up to Grade V, while Tsang et al. [16] sampled up to Grade III).

Microbial communities can assemble through stochastic or deterministic assembly processes. Stochastic processes include priority effects and ecological drift, while deterministic processes include more selective factors such as species traits, interspecies interactions (e.g., competition, mutualism), and environmental filtering. The observation of a host phylogeny component to microbial community patterns suggests the presence of host selection on the gut community, whether directly through host immune responses, or indirectly through feeding preferences or shared behaviors. This tight coevolution between a host and its microbial community is referred to as phylosymbiosis [68, 69]. While previous work in marine invertebrates found no evidence of large-scale phylosymbiosis [70], Tsang et al. [16] reported that the variation in the gut microbiome explained by mangrove crab host identity decreased at higher taxonomic groupings. This finding suggests that phylosymbiosis may be occurring at more recent (i.e. genus-level) evolutionary scales in crab lineages, which largely aligns with our results from crabs with greater degrees of terrestrial adaptation.

### Relative abundances of specific taxa vary across diet, terrestrial grade, and host genus

At the phylum level, our findings for bacterial community relative abundance patterns are broadly consistent with previous studies in crabs and other crustaceans [16, 17, 71, 72], where Proteobacteria (or Pseudomonadota) dominate, followed by Bacteroidota (Bacteroidetes), Bacillota (Firmicutes), and Actinomycetota (Actinobacteria). However, the proportions reported here slightly differ from other studies, which may reflect ecological or methodological differences. For instance, studies of mangrove crabs report relative abundances of Proteobacteria ranging from 44.3% to 86.1% and Bacteroidetes from 11.1% to 34.6% [71, 72], while our findings using K2/B and MAG-based analyses show relative abundances of 54.4% and 34.5%, respectively for Proteobacteria, and 9.86% and 44.8% respectively for Bacteroidota. Since many of the crab species used in our study display relatively greater degrees of terrestrial adaptation than those used in previous work, differences in the specific ecologies of the crabs may be driving these differences in bacterial community relative abundance patterns. Future studies using crabs across the full gradient of terrestriality may further illuminate these patterns.

At finer taxonomic levels, several bacterial genera, such as *Citrobacter* and *Paracoccus*, exhibited significant shifts in abundance across gradients of terrestriality, diet, and host genus. Notably, members of *Citrobacter* and *Paracoccus* can be lignin degraders [73–76], and these groups were more abundant in crabs from higher terrestrial grades. In combination with the broad scale functional results, where lower grade crabs lacked complete complex carbon degradation pathways compared to higher grade crabs, this suggests that there may be more selection for microbes with lignin degradation and other complex carbon degradation pathways at higher terrestrial grades.

Few studies have explored the diversity and role of archaea and fungi in crab gut microbiomes. Here, uncultured and undescribed lineages dominated both groups, consistent with their representation as rare yet potentially functionally important members of gut communities. For fungi, *Trametes* exhibited significant variation in relative abundance across terrestrial grades, but generally was more prevalent in crabs from higher terrestrial grades. This aligns with reports of *Trametes* in insect gut microbiomes, where it degrades structural components of lignocellulose [77]. Archaea were generally less abundant across all collected samples, but we did observe some significant differences in relative abundance across diet (*Methanobrevibacter* was more abundant in carnivores, while *Natrinema* was more abundant in detritivores and herbivores) and host genus (unclassified *Candidatus* Methanoperedenaceae was found only in *Birgus*). Archaea and fungi both may have important functional roles in terrestrial animal guts including in lignocellulose degradation [14]; future work should use a more targeted approach (i.e., amplicon sequencing) to better characterize these communities and their functional roles in the guts of terrestrial crabs.

### Functional convergence across crab gut microbiomes

We observed broad scale differences in the functional potential of the gut microbiomes of crabs across terrestrial grades, potentially reflecting adaptations to dietary and environmental shifts. We observed broad patterns supporting complex carbon pathway enrichment in the microbiomes of crabs from higher terrestrial grades, contrasting with the more limited presence of these pathways in crabs from lower grades. This gradient aligns with the increasing reliance on terrestrial plant material as a dietary component in more terrestrially adapted species, supporting the hypothesis that gut microbial assembly and function are strongly influenced by diet and habitat.

We found that presence of CAZyme domains related to lignocellulose degradation varied across bacterial classes and host crab species. Generally, we observed glycoside hydrolases with potential roles in cellulose and hemicellulose breakdown (GH2, GH3, GH5, GH16, GH20, GH29, GH30), alpha-glucan-active GH13, lignin-oxidizing enzymes (AA1, AA3), lignin-degrading auxiliary enzymes (AA4), and carbohydrate esterases (CE1, CE4). GH2 was the most abundant lignocellulose-related family, and GH3 and GH13 were the most widely distributed. GH13s were also the most prevalent CAZyme families detected in metagenomes from terrestrial and aquatic isopod guts, suggesting that these likely play a critical role in arthropod gut metabolism [78].

Microbial GH1, GH5, and GH30 have been implicated in lignocellulose degradation in a variety of animal hosts [14], and were observed here in MAGs in the classes Bacteroidia, Clostridia and Gammaproteobacteria (phylum Bacteroidota, Firmicutes, and Pseudomonadota respectively). Similar to our results, GH1, GH2, GH3, and AA1 were previously found in higher abundance in herbivorous and detritivorous crab metagenomes, with contributions largely from Pseudomonadota and Bacteroidota [17]. In our study, Bacteroidota was the phyla with the largest number of recovered high-quality MAGs, highest metagenomic prevalence (44.80%), and largest diversity of CAZyme domains associated with lignocellulose degradation related functions including GH2, GH3, GH5, and GH30. Bacteroidota lineages dominate many terrestrial lignocellulose-degrading communities [79], and are also keystone taxa in the gut microbiomes of various animals, including humans [80].

It is clear that the gut microbiomes of crabs encode a diverse set of cellulolytic enzymes, with even non-herbivorous crabs hosting microbes with at least some genetic potential for this activity, as has previously been reported in more aquatic crabs [17]. Culture based studies have previously highlighted potential roles for *Demequina* and *Paenibacillus* in cellulose degradation in the intestines of more aquatic crabs [71, 81]. These studies found these bacteria both in crab guts, but also in their local habitat (e.g., burrow walls) suggesting that they were being environmentally acquired. While we recovered no MAGs belonging to *Paenibacillus*, MAGs from *Demequina* were enriched in several CAZyme domains including lignocellulose degradation enzymes (GH1, GH3) and lignin-oxidizing enzymes (AA1, AA3). One reason that might explain why we observe lignocellulose degrading CAZyme domains in the gut microbiomes of crabs from lower grades of terrestriality is the importance of maintaining limited carbon degradation functioning due to their diet. For example, aquatic crab diets can contain algae, seagrasses, and other marine detritus which may have low levels of lignocellulose and other complex carbon compounds [10]. However, the diets of higher terrestrial grade crabs contain a much larger proportion of lignocellulose containing food [11], consistent with the observed enrichment of known lignin degrading microbes and broad-scale functional pathways related to these functions in higher grade crabs. Overall, these functional results, coupled with the importance of habitat and diet to overall microbial community composition, support a hypothesis of a mechanism of horizontal or environmental acquisition of microbes with lignocellulose degradation functions by crabs. Future studies should investigate crab gut community assembly through source tracking (e.g., by sampling local environment, diets, etc.) or through laboratory experiments controlling for specific diet and environmental challenges. Additionally, research should be performed to rule out vertical transmission through studies aimed at parental manipulation, antibiotic treatment of eggs, or combinatorial treatments.

### Study limitations

Several sampling and analytical limitations should be considered when interpreting our findings. As many of the animals we sampled for this work were individuals from protected species collected from national parks, we conducted this study while operating within the constraints of our permits, which restricted our collection efforts. Our limited total sample sizes (n = 32 individuals across 14 species and 6 genera) led to representational imbalances across genera, diet, and terrestrial grades. Specifically, while herbivores (n = 13) and detritivores (n = 13) were well-represented, omnivores (n = 5) and carnivores (n = 1) had limited replication. Furthermore, as host phylogeny, terrestrial grade, and geographic origin partially covaried across our samples, some microbial variation attributed to host genus or degree of terrestrialization may also reflect regional environmental pools or other unexamined habitat differences.

Additionally, sample size constraints directly impact the interpretation of our Random Forest analysis. Although we achieved high accuracy overall (85.71%) using complex carbon degradation gene counts, a 30% hold-out test set is composed of only 9 samples. Our performance metrics are therefore inherently sensitive to the classification of individual samples, especially within our undersampled categories (e.g. omnivores and carnivores) where our predictive accuracy was the lowest. Our findings from our Random Forest model, while generally corroborating our broader functional conclusions, should therefore be interpreted as exploratory evidence for diet-associated divergences in gut microbial functions.

Finally, because shotgun metagenomics characterizes genetic potential rather than active gene expression or enzymatic rates, the presence of predicted lignocellulose-degrading CAZyme domains does not inherently confirm these metabolic activities within the crab gut. Future studies incorporating larger, more balanced sampling should also pair metagenomic approaches with metatranscriptomics, metabolomics, and controlled feeding experiments to further disentangle the effects of host phylogeny, diet, and environmental or vertical microbial acquisition on convergent terrestrial crab herbivory.

## Conclusion

This study provides a comprehensive analysis of land crab gut microbial communities across a robust terrestrial gradient using metagenomics. We constructed a high-quality genomic resource of 129 MAGs representing dominant bacterial members, offering a valuable foundation for understanding crab-bacterial interactions and evolution, with the aim of interrogating what role, if any, particular gut microbial community assemblies may have played in the terrestrialization of the land crabs. Our findings show that host crab genus is the primary driver of bacterial community composition, with the degree of host crab terrestriality also significantly contributing. Multiple bacterial genera varied in relative abundance across host terrestrial grades, taxonomic affinity, and diets, highlighting the dynamic nature of these microbial communities. Broad-scale analysis of carbon metabolism revealed the absence of key carbon degradation metabolic pathways (carbon-in crabs from lower terrestrial grades, while fine-scale functional analyses linked specific MAGs to CAZyme domains reported to be involved in lignocellulose-degradation pathways. These results suggest that different crab genera may rely on distinct microbial taxa to perform similar functions, demonstrating possible convergent acquisition of microbes with shared functional roles to support lignocellulose degradation. Future research should investigate the mechanisms of microbial acquisition, whether through environmental exposure or vertical transmission, which may allow us to further reconstruct the evolutionary histories of these essential gut symbioses and determine more precisely whether selective or phylogenetic factors are critical drivers of the observed patterns of association.

## Declarations

Ethics approval and consent to participate:

Not applicable

## Consent for publication

Not applicable

## Availability of data and material

This metagenomic sequencing project and raw metagenomic sequencing reads have been deposited at GenBank under BioProject PRJNA1141481, at the accession no. SRR30067231-SRR30067263. MAG assemblies are also available at NCBI under accession no. JBKDZH000000000-JBKEEF000000000. Mitogenomes are archived on Zenodo (doi: 10.5281/zenodo.21459538). All data and code can be found in GitHub (https://github.com/casett/LandCrabMicrobiomes) and is archived on Zenodo (doi: 10.5281/zenodo.14895641).

## Authors’ contributions

C.L.E and V.W.Z conceived and designed the experiments and obtained funding. V.W.Z and L.G.E.W. performed field collections in Australia and Panamá, respectively. A.H.contributed with Panamanian field collections and species identifications. C.L.E., V.W.Z, and L.G.E.W extracted DNA, analyzed the data, prepared figures and/or tables, and wrote the paper. R.K.G. and J.A.E. provided advice on experimental design. All authors reviewed drafts of the paper.

## Acknowledgements

We would like to thank members of the UC Davis Genome Center DNA Technologies Core (https://dnatech.genomecenter.ucdavis.edu/) for preparing libraries and performing sequencing of metagenomic samples. We would also like to thank the park rangers at the Christmas Island National Park for their field collection assistance, and all of the land crabs that were sacrificed in service of this work. In Panamá, we would like to thank the staff at the Bocas del Toro Research Station of the Smithsonian Tropical Research Institute who hosted us during our field trip and granted access to their laboratories.

## References

1. Grosberg RK, Vermeij GJ, Wainwright PC. Biodiversity in water and on land. Curr Biol. 2012;22.

2. Watson-Zink VM. Making the grade: Physiological adaptations to terrestrial environments in decapod crabs. Arthropod Struct Dev. 2021;64:101089.

3. Vermeij GJ, Dudley R. Why are there so few evolutionary transitions between aquatic and terrestrial ecosystems? Biol J Linn Soc Lond. 2008;70:541–54.

4. Sues H-D, editor. Evolution of Herbivory in Terrestrial Vertebrates: Perspectives from the Fossil Record. Cambridge University Press; 2000.

5. Kergoat GJ, Meseguer AS, Jousselin E. Evolution of Plant–Insect Interactions: Insights From Macroevolutionary Approaches in Plants and Herbivorous Insects. In: Advances in Botanical Research. Academic Press; 2017. p. 25–53.

6. Bredon M, Herran B, Lheraud B, Bertaux J, Grève P, Moumen B, et al. Lignocellulose degradation in isopods: new insights into the adaptation to terrestrial life. BMC Genomics. 2019;20.

7. Ord TJ, Hundt PJ. Crossing extreme habitat boundaries: Jack-of-all-trades facilitates invasion but is eroded by adaptation to a master-of-one. Funct Ecol. 2020;34:1404–15.

8. Wang Z, Xu S, Du K, Huang F, Chen Z, Zhou K, et al. Evolution of Digestive Enzymes and RNASE1 Provides Insights into Dietary Switch of Cetaceans. Mol Biol Evol. 2016;33:3144–57.

9. Wolfe JM, Ballou L, Luque J, Watson-Zink VM, Ahyong ST, Barido-Sottani J, et al. Convergent Adaptation of True Crabs (Decapoda: Brachyura) to a Gradient of Terrestrial Environments. Syst Biol. 2024;73:247–62.

10. Ropes JW. The Feeding Habits of the Green Crab, Carcinus Maenas (L.). Fishery Bulletin. 1968;67:183–203.

11. Greenaway P, Raghaven S. Digestive strategies in two species of leaf-eating land crabs (Brachyura: Gecarcinidae) in a rain forest. Physiol Zool. 1998;71.

12. Allardyce BJ, Linton SM, Saborowski R. The last piece in the cellulase puzzle: the characterisation of beta-glucosidase from the herbivorous gecarcinid land crab *Gecarcoidea natalis*. J Exp Biol. 2010;213 Pt 17.

13. Ruiz-Dueñas FJ, Martínez AT. Microbial degradation of lignin: how a bulky recalcitrant polymer is efficiently recycled in nature and how we can take advantage of this. Microb Biotechnol. 2009;2:164–77.

14. Cragg SM, Beckham GT, Bruce NC, Bugg TD, Distel DL, Dupree P, et al. Lignocellulose degradation mechanisms across the Tree of Life. Curr Opin Chem Biol. 2015;29.

15. Hsu C-H, Soong K. Mechanisms causing size differences of the land hermit crab *Coenobita rugosus* among eco-islands in Southern Taiwan. PLoS One. 2017;12.

16. Tsang CTT, Hui TKL, Chung NM, Yuen WT, Tsang LM. Comparative analysis of gut microbiome of mangrove brachyuran crabs revealed patterns of phylosymbiosis and codiversification. Molecular ecology. 2024;33.

17. Hui TKL, Lo ICN, Wong KKW, Tsang CTT, Tsang LM. Metagenomic analysis of gut microbiome illuminates the mechanisms and evolution of lignocellulose degradation in mangrove herbivorous crabs. BMC microbiology. 2024;24.

18. Cannicci S, Fratini S, Meriggi N, Bacci G, Iannucci A, Mengoni A, et al. To the Land and Beyond: Crab Microbiomes as a Paradigm for the Evolution of Terrestrialization. Front Microbiol. 2020;11.

19. Meng G, Li Y, Yang C, Liu S. MitoZ: a toolkit for animal mitochondrial genome assembly, annotation and visualization. Nucleic Acids Res. 2019;47:e63.

20. Tan MH, Gan HM, Schultz MB, Austin CM. MitoPhAST, a new automated mitogenomic phylogeny tool in the post-genomic era with a case study of 89 decapod mitogenomes including eight new freshwater crayfish mitogenomes. Mol Phylogenet Evol. 2015;85:180–8.

21. Bushnell B. BBMap: a fast, accurate, splice-aware aligner. Lawrence Berkeley National Lab.(LBNL), Berkeley, CA (United States); 2014.

22. Wood DE, Lu J, Langmead B. Improved metagenomic analysis with Kraken 2. Genome Biol. 2019;20:257.

23. Lu J, Breitwieser FP, Thielen P, Salzberg SL. Bracken: estimating species abundance in metagenomics data. PeerJ Comput Sci. 2017;3:e104.

24. Huerta-Cepas J, Serra F, Bork P. ETE 3: Reconstruction, Analysis, and Visualization of Phylogenomic Data. Mol Biol Evol. 2016;33:1635–8.

25. Li D, Liu C-M, Luo R, Sadakane K, Lam T-W. MEGAHIT: an ultra-fast single-node solution for large and complex metagenomics assembly via succinct de Bruijn graph. Bioinformatics. 2015;31:1674–6.

26. Kang DD, Li F, Kirton E, Thomas A, Egan R, An H, et al. MetaBAT 2: an adaptive binning algorithm for robust and efficient genome reconstruction from metagenome assemblies. PeerJ. 2019;7:e7359.

27. Wu Y-W, Tang Y-H, Tringe SG, Simmons BA, Singer SW. MaxBin: an automated binning method to recover individual genomes from metagenomes using an expectation-maximization algorithm. Microbiome. 2014;2:26.

28. Graham ED, Heidelberg JF, Tully BJ. BinSanity: unsupervised clustering of environmental microbial assemblies using coverage and affinity propagation. PeerJ. 2017;5:e3035.

29. Alneberg J, Bjarnason BS, de Bruijn I, Schirmer M, Quick J, Ijaz UZ, et al. Binning metagenomic contigs by coverage and composition. Nat Methods. 2014;11:1144–6.

30. Pronk LJU, Medema MH. Whokaryote: distinguishing eukaryotic and prokaryotic contigs in metagenomes based on gene structure. Microbial genomics. 2022;8.

31. Sieber CMK, Probst AJ, Sharrar A, Thomas BC, Hess M, Tringe SG, et al. Recovery of genomes from metagenomes via a dereplication, aggregation and scoring strategy. Nat Microbiol. 2018;3:836–43.

32. Chklovski A, Parks DH, Woodcroft BJ, Tyson GW. CheckM2: a rapid, scalable and accurate tool for assessing microbial genome quality using machine learning. bioRxiv. 2022.

33. Reiter TE, Tessa Pierce-Ward N, Irber L, Schwarz EM, Titus Brown C. Charcoal: filtering contamination in metagenome-assembled genome bins and other genomes. Manubot. 2022.

34. Nayfach S, Shi ZJ, Seshadri R, Pollard KS, Kyrpides NC. New insights from uncultivated genomes of the global human gut microbiome. Nature. 2019;568:505–10.

35. Bowers RM, Kyrpides NC, Stepanauskas R, Harmon-Smith M, Doud D, Reddy TBK, et al. Minimum information about a single amplified genome (MISAG) and a metagenome-assembled genome (MIMAG) of bacteria and archaea. Nat Biotechnol. 2017;35:725–31.

36. Chaumeil P-A, Mussig AJ, Hugenholtz P, Parks DH. GTDB-Tk: a toolkit to classify genomes with the Genome Taxonomy Database. Bioinformatics. 2019. 10.1093/bioinformatics/btz848.

37. Aroney STN, Newell RJP, Nissen J, Camargo AP, Tyson GW, Woodcroft BJ. CoverM: Read coverage calculator for metagenomics. 2024. 10.5281/zenodo.10531254.

38. Li H. Minimap2: pairwise alignment for nucleotide sequences. Bioinformatics. 2018;34:3094–100.

39. The R Project for Statistical Computing. https://www.r-project.org/.

40. Wickham H, Averick M, Bryan J, Chang W, McGowan L, François R, et al. Welcome to the Tidyverse. Journal of Open Source Software. 2019;4:1686.

41. McMurdie PJ, Holmes S. phyloseq: An R Package for Reproducible Interactive Analysis and Graphics of Microbiome Census Data. PLoS ONE. 2013;8:e61217.

42. Oksanen J, Simpson G, Blanchet F, Kindt R, Legendre P, Minchin P, et al. vegan: Community Ecology Package. 2024.

43. Davis NM, Proctor DM, Holmes SP, Relman DA, Callahan BJ. Simple statistical identification and removal of contaminant sequences in marker-gene and metagenomics data. Microbiome. 2018;6:226.

44. Martinez Arbizu P. pairwiseAdonis: Pairwise multilevel comparison using adonis. 2020.

45. Shaffer M, Borton MA, McGivern BB, Zayed AA, La Rosa SL, Solden LM, et al. DRAM for distilling microbial metabolism to automate the curation of microbiome function. Nucleic Acids Res. 2020;48:8883–900.

46. Hyatt D, Chen G-L, Locascio PF, Land ML, Larimer FW, Hauser LJ. Prodigal: prokaryotic gene recognition and translation initiation site identification. BMC Bioinformatics. 2010;11:119.

47. Aramaki T, Blanc-Mathieu R, Endo H, Ohkubo K, Kanehisa M, Goto S, et al. KofamKOALA: KEGG Ortholog assignment based on profile HMM and adaptive score threshold. Bioinformatics. 2020;36:2251–2.

48. Finn RD, Bateman A, Clements J, Coggill P, Eberhardt RY, Eddy SR, et al. Pfam: the protein families database. Nucleic Acids Res. 2014;42 Database issue:D222–30.

49. Lombard V, Golaconda Ramulu H, Drula E, Coutinho PM, Henrissat B. The carbohydrate-active enzymes database (CAZy) in 2013. Nucleic Acids Res. 2014;42 Database issue:D490–5.

50. Huang L, Zhang H, Wu P, Entwistle S, Li X, Yohe T, et al. dbCAN-seq: a database of carbohydrate-active enzyme (CAZyme) sequence and annotation. Nucleic Acids Res. 2018;46:D516–21.

51. Rawlings ND, Barrett AJ, Bateman A. Using the MEROPS Database for Proteolytic Enzymes and Their Inhibitors and Substrates. Curr Protoc Bioinformatics. 2014;48:1.25.1–33.

52. Janusz G, Pawlik A, Sulej J, Swiderska-Burek U, Jarosz-Wilkolazka A, Paszczynski A. Lignin degradation: microorganisms, enzymes involved, genomes analysis and evolution. FEMS Microbiol Rev. 2017;41:941–62.

53. Levasseur A, Drula E, Lombard V, Coutinho PM, Henrissat B. Expansion of the enzymatic repertoire of the CAZy database to integrate auxiliary redox enzymes. Biotechnol Biofuels. 2013;6:41.

54. Liu Y, Wu Y, Zhang Y, Yang X, Yang E, Xu H, et al. Lignin degradation potential and draft genome sequence of S0301. Biotechnol Biofuels. 2019;12:256.

55. Chettri D, Nad S, Konar U, Verma AK. CAZyme from gut microbiome for efficient lignocellulose degradation and biofuel production. Front Chem Eng Chin. 2022;4:1054242.

56. Sabbadin F, Pesante G, Elias L, Besser K, Li Y, Steele-King C, et al. Uncovering the molecular mechanisms of lignocellulose digestion in shipworms. Biotechnol Biofuels. 2018;11:59.

57. King AJ, Cragg SM, Li Y, Dymond J, Guille MJ, Bowles DJ, et al. Molecular insight into lignocellulose digestion by a marine isopod in the absence of gut microbes. Proc Natl Acad Sci U S A. 2010;107:5345–50.

58. Kolde R. pheatmap: Pretty Heatmaps. 2018.

59. Bankevich A, Nurk S, Antipov D, Gurevich AA, Dvorkin M, Kulikov AS, et al. SPAdes: a new genome assembly algorithm and its applications to single-cell sequencing. J Comput Biol. 2012;19:455–77.

60. Zhou Z, Tran PQ, Breister AM, Liu Y, Kieft K, Cowley ES, et al. METABOLIC: high-throughput profiling of microbial genomes for functional traits, metabolism, biogeochemistry, and community-scale functional networks. Microbiome. 2022;10:1–22.

61. Wright MN, Ziegler A. Ranger: A fast implementation of random forests for high dimensional data in C++ and R. J Stat Softw. 2017;77.

62. Kuhn M. Building Predictive Models in R Using the caret Package. J Stat Softw. 2008;28.

63. Balvočiūtė M, Huson DH. Silva, RDP, Greengenes, NCBI and OTT — how do these taxonomies compare? BMC Genomics. 2017;18:1–8.

64. Parks DH, Chuvochina M, Waite DW, Rinke C, Skarshewski A, Chaumeil P-A, et al. A standardized bacterial taxonomy based on genome phylogeny substantially revises the tree of life. Nat Biotechnol. 2018;36:996–1004.

65. Portik DM, Brown CT, Pierce-Ward NT. Evaluation of taxonomic classification and profiling methods for long-read shotgun metagenomic sequencing datasets. BMC Bioinformatics. 2022;23:1–39.

66. Herlemann DPR, Tammert H, Kivistik C, Käiro K, Kisand V. Distinct biogeographical patterns in snail gastrointestinal tract bacterial communities compared with sediment and water. Microbiologyopen. 2024;13:e13.

67. Hu Z, Tong Q, Chang J, Xu J, Wu B, Han Y, et al. Host species of freshwater snails within the same freshwater ecosystem shapes the intestinal microbiome. Front Ecol Evol. 2024;12:1341359.

68. Lim SJ, Bordenstein SR. An introduction to phylosymbiosis. Proceedings Biological sciences. 2020;287.

69. Brooks AW, Kohl KD, Brucker RM, van Opstal EJ, Bordenstein SR. Phylosymbiosis: Relationships and Functional Effects of Microbial Communities across Host Evolutionary History. PLoS biology. 2016;14.

70. Boscaro V, Holt CC, Van Steenkiste NWL, Herranz M, Irwin NAT, Àlvarez-Campos P, et al. Microbiomes of microscopic marine invertebrates do not reveal signatures of phylosymbiosis. Nature microbiology. 2022;7.

71. Tongununui P, Kuriya Y, Murata M, Sawada H, Araki M, Nomura M, et al. Mangrove crab intestine and habitat sediment microbiomes cooperatively work on carbon and nitrogen cycling. PloS one. 2021;16.

72. Shao C, Zhao W, Li N, Li Y, Zhang H, Li J, et al. Gut Microbiome Succession in Chinese Mitten Crab During Seawater-Freshwater Migration. Front Microbiol. 2022;13:858508.

73. Cortes-Tolalpa L, Wang Y, Salles JF, van Elsas JD. Comparative Genome Analysis of the Lignocellulose Degrading Bacteria so4 and w15. Front Microbiol. 2020;11:248.

74. Pyter W, Grewal J, Bartosik D, Drewniak L, Pranaw K. Pigment Production by *Paracoccus* spp. Strains through Submerged Fermentation of Valorized Lignocellulosic Wastes. Fermentation. 2022;8:440.

75. Sharker B, Islam MA, Hossain MAA, Ahmad I, Al Mamun A, Ghosh S, et al. Characterization of lignin and hemicellulose degrading bacteria isolated from cow rumen and forest soil: Unveiling a novel enzymatic model for rice straw deconstruction. Sci Total Environ. 2023;904:166704.

76. Asina F, Brzonova I, Voeller K, Kozliak E, Kubátová A, Yao B, et al. Biodegradation of lignin by fungi, bacteria and laccases. Bioresource technology. 2016;220.

77. Rojas-Jiménez K, Hernández M. Isolation of Fungi and Bacteria Associated with the Guts of Tropical Wood-Feeding Coleoptera and Determination of Their Lignocellulolytic Activities. International journal of microbiology. 2015;2015.

78. Bredon M, Herran B, Bertaux J, Grève P, Moumen B, Bouchon D. Isopod holobionts as promising models for lignocellulose degradation. Biotechnol Biofuels. 2020;13:49.

79. Naas AE, Mackenzie AK, Mravec J, Schückel J, Willats WGT, Eijsink VGH, et al. Do rumen Bacteroidetes utilize an alternative mechanism for cellulose degradation? mBio. 2014;5:e01401–14.

80. Shin JH, Tillotson G, MacKenzie TN, Warren CA, Wexler HM, Goldstein EJC. Bacteroides and related species: The keystone taxa of the human gut microbiota. Anaerobe. 2024;85:102819.

81. Lee CY, Lee SY. Contribution of Aerobic Cellulolytic Gut Bacteria to Cellulose Digestion in Fifteen Coastal Grapsoid Crabs Underpins Potential for Mineralization of Mangrove Production. Curr Microbiol. 2024;81:224.

