## Supplementary Materials for "Convergent lignocellulose degradation in terrestrial crabs is driven by distinct host genus-specific microbial communities"

### Supplemental Materials

**Figure S1. Bacterial, archaeal and fungal beta diversity across diet, terrestrial grades and crab genera.** Principal-coordinate analysis (PCoA) visualization of Bray Curtis dissimilarities of Kraken2/Bracken counts of bacterial abundance showing individual samples designated as points with colors and shapes representing (A) diet, (B) terrestrial grade, and (C) crab genera. Archaeal abundance of (D) diet, (E) terrestrial grade, and (F) crab genera, and fungal abundance of (G) diet, (H) terrestrial grade, and (I) crab genera.

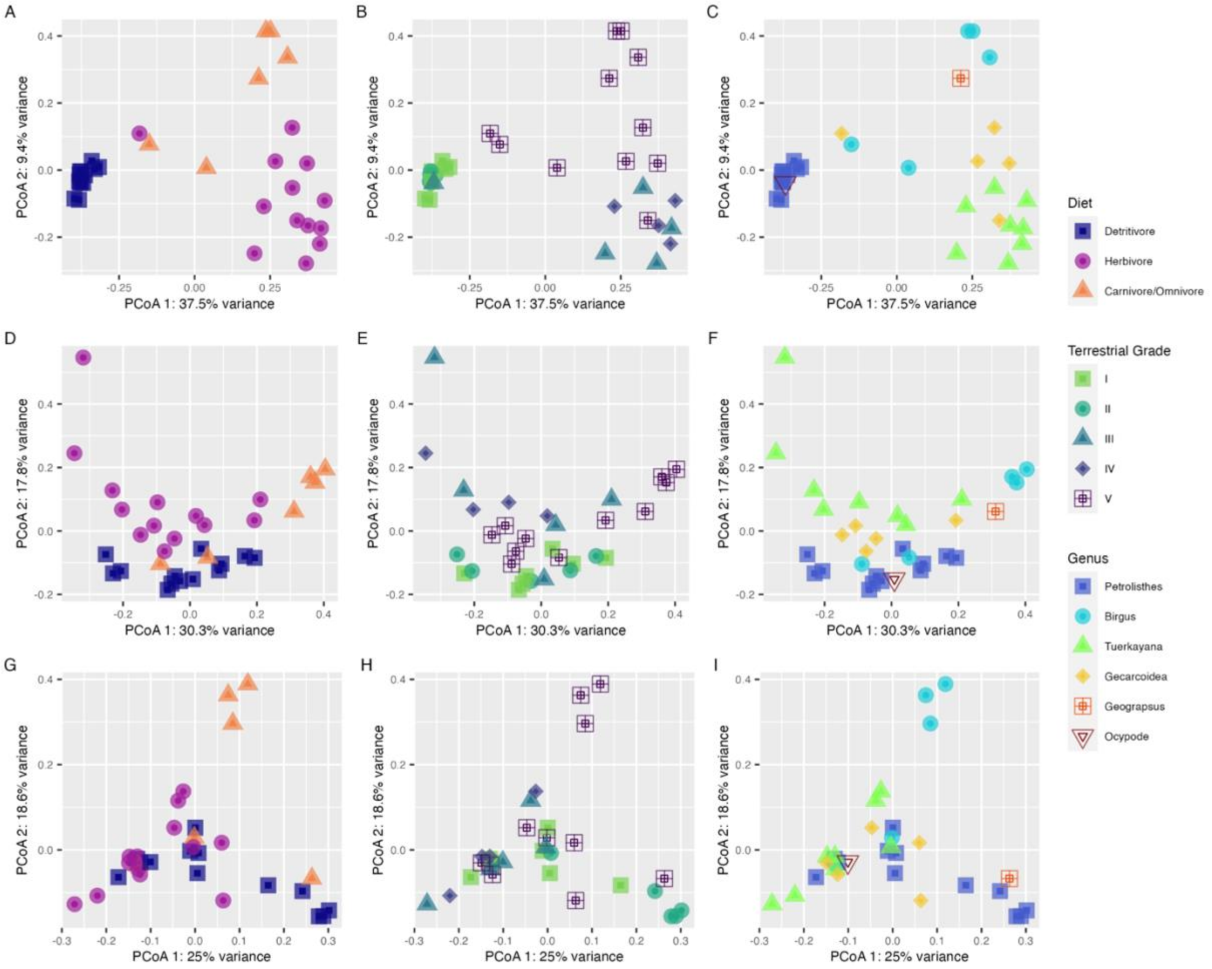

**Figure S2. Mean relative abundance of bacterial classes associated with crab species.** Stacked bar charts displaying the mean relative abundance, colored by predicted taxonomic class, of bacteria identified by Kraken2/Bracken (left) and bacterial MAGs estimated using coverM (right). Classes representing less than 5% mean relative abundance across either dataset are collapsed for visualization purposes into a single group labeled ‘Other’.

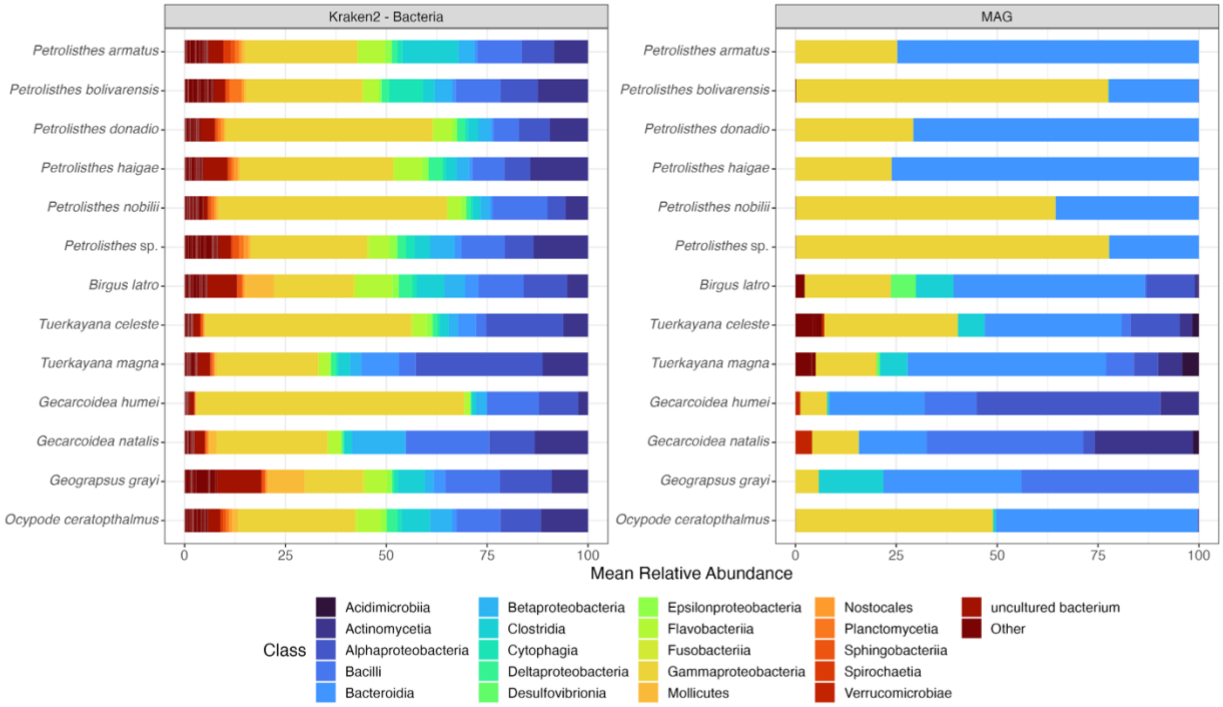

**Figure S3. Mean relative abundance of archaeal classes associated with crab species.** Stacked bar charts displaying the mean relative abundance, colored by predicted class, of archaea identified by Kraken2/Bracken. Classes representing less than 5% mean relative abundance across the dataset are collapsed for visualization purposes into a single group labeled 'Other'.

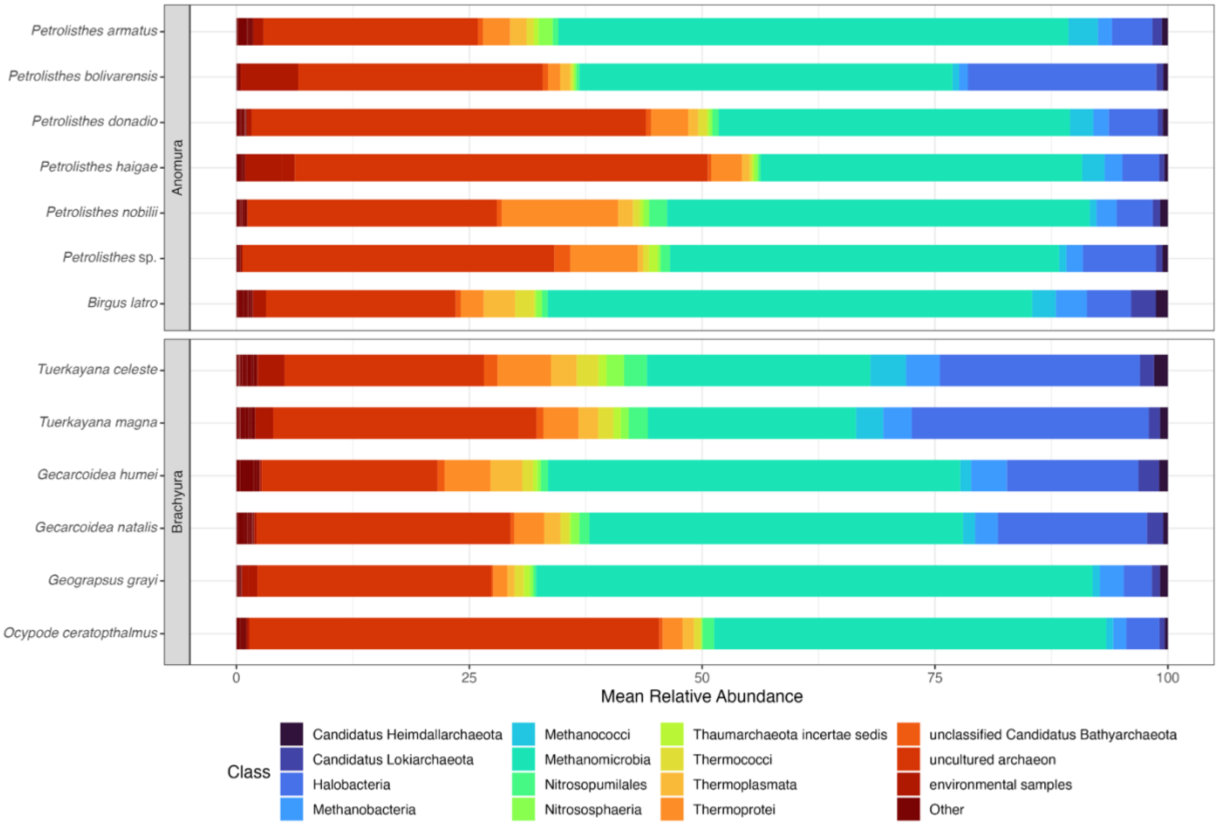

**Figure S4. Mean relative abundance of fungal classes associated with crab species.** Stacked bar charts displaying the mean relative abundance, colored by predicted class, of fungi identified by Kraken2/Bracken. Classes representing less than 5% mean relative abundance across the dataset are collapsed for visualization purposes into a single group labeled 'Other'.

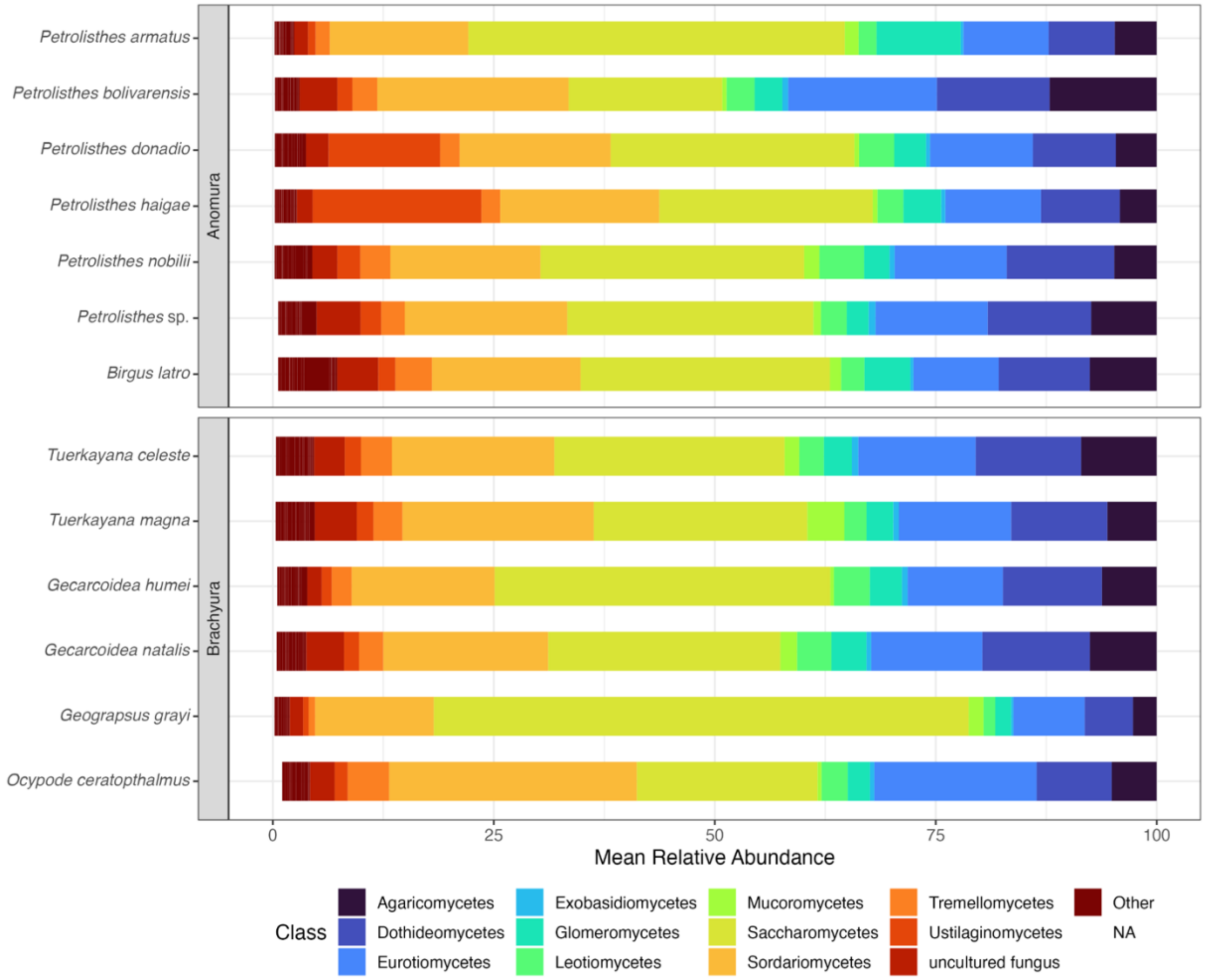

**Figure S5. Microbial genera whose relative abundance differs significantly across diets, grades and host genera.** Bar charts display the mean relative abundance of (A) bacterial and (B) fungal genera identified by Kraken2/Bracken whose relative abundance differed significantly with crab diet, terrestrial grade and genera. Bars are colored by predicted genus and error bars represent standard error.

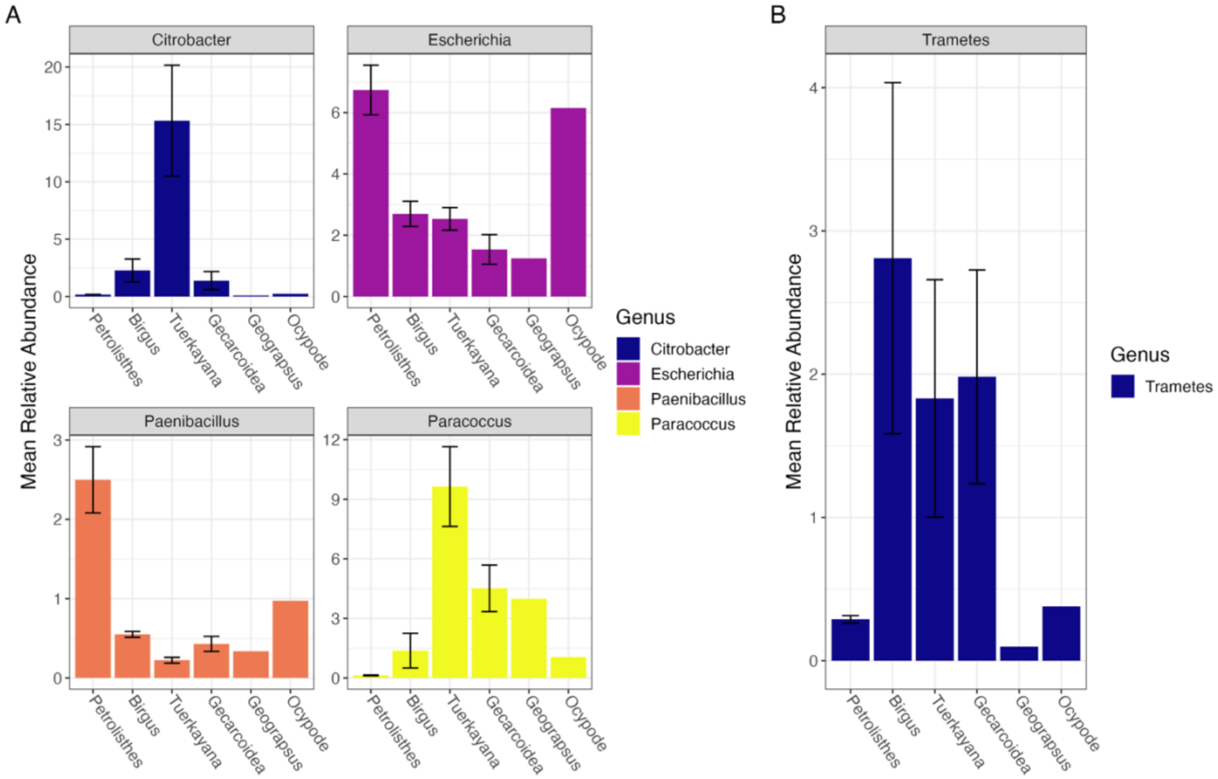

**Figure S6. CAZyme domain abundance across MAGs. DRAM was used to annotate CAZyme domains in the proteome of MAGs.** A heatmap displaying the log-transformed counts of CAZyme domains with potential lignocellulose-related functions across all MAGs. Specific CAZyme modules are denoted across the top, while MAG taxonomic order and class information are denoted on the left. Further, the dendrogram to the left of the heatmap clusters the MAGs by similarity in counts between the different CAZyme domains, while the dendrogram above clusters CAZymes based on similarity in counts between the different MAGs.

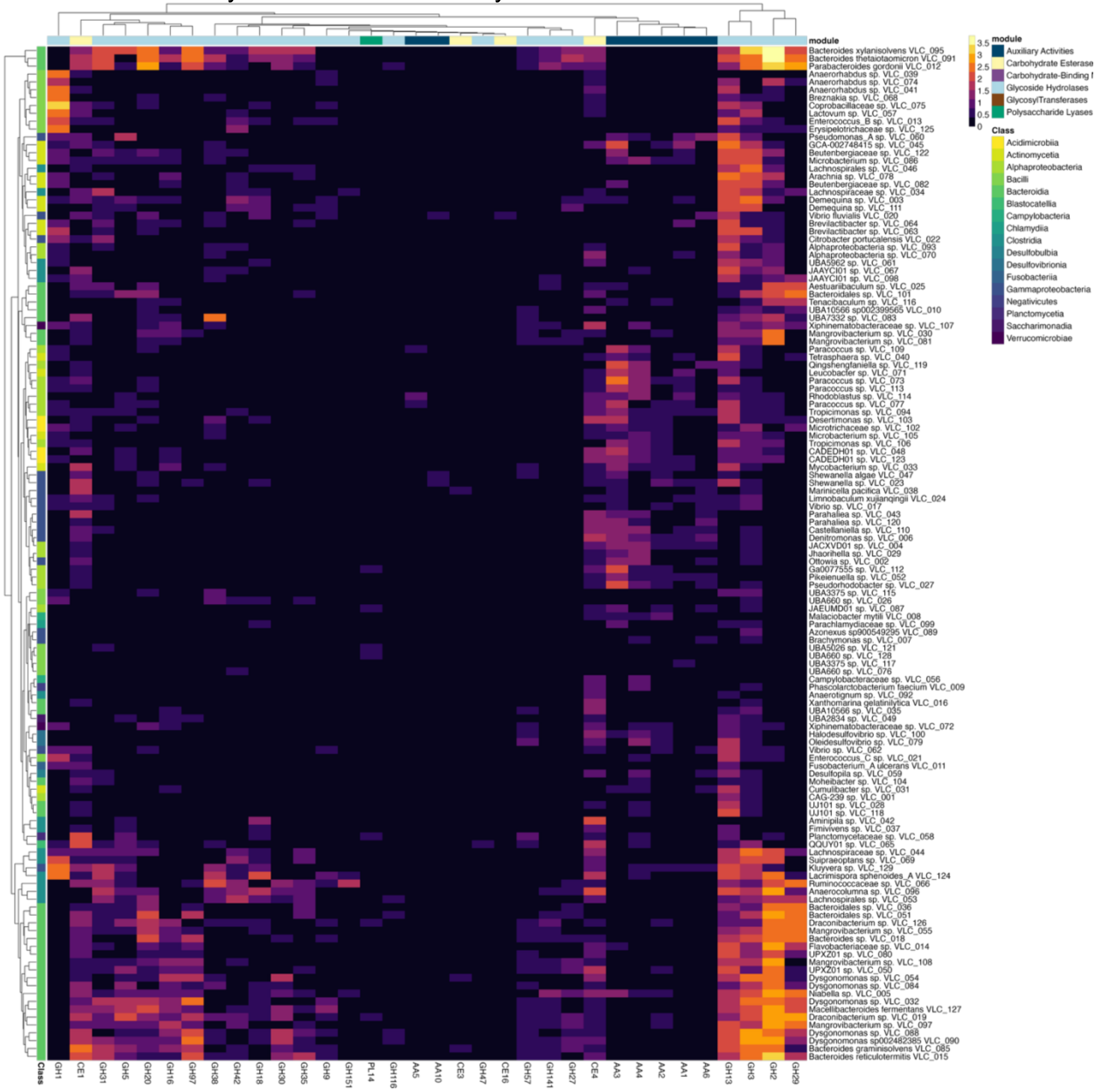

**Figure S7. CAZyme domain abundance across Clostridia.** (A) DRAM was used to annotate CAZyme domains in the proteome of MAGs. A heatmap displaying the log-transformed counts of CAZyme domains with potential lignocellulose-related functions for members of the Clostridia. Specific CAZyme modules are denoted across the top, while MAG taxonomic order and class information are denoted on the left.  Further, the dendrogram to the left of the heatmap clusters the MAGs by similarity in counts between the different CAZyme domains, while the dendrogram above clusters CAZymes based on similarity in counts between the different MAGs. (B) Relative abundance of MAGs in the Clostridia across host crab genera are displayed as a barplot. Bars are split by taxonomic genus, colored by predicted order and error bars represent standard error.

(A)

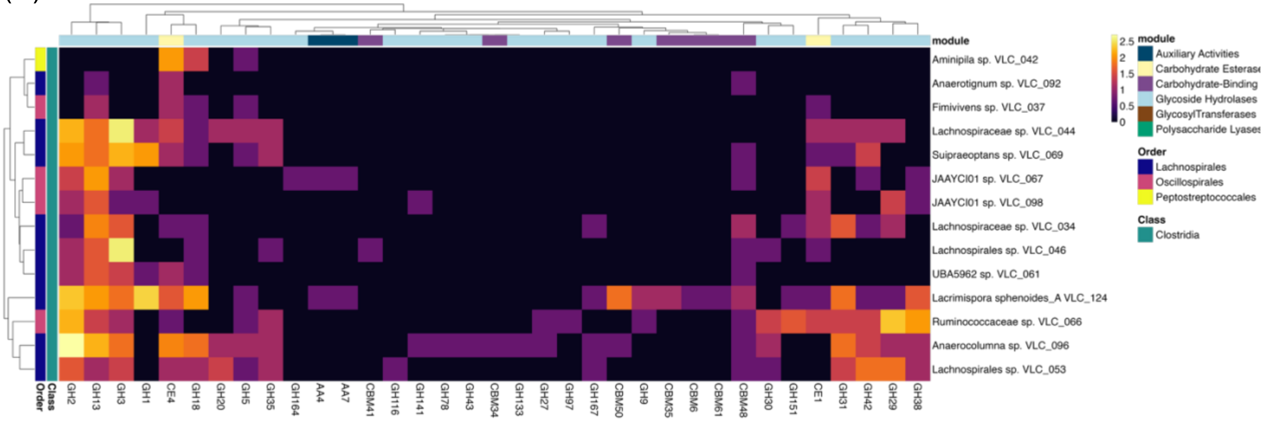


(B)

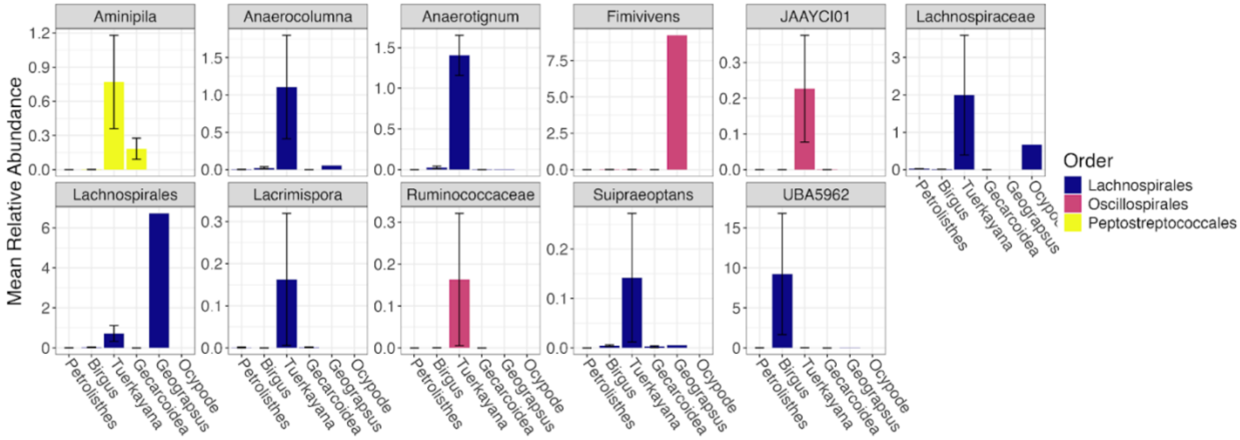

**Figure S8. CAZyme domain abundance across Alphaproteobacteria.** (A) DRAM was used to annotate CAZyme domains in the proteome of MAGs. A heatmap displaying the log-transformed counts of CAZyme domains with potential lignocellulose-related functions for members of the Alphaproteobacteria. Specific CAZyme modules are denoted across the top, while MAG taxonomic order and class information are denoted on the left. Further, the dendrogram to the left of the heatmap clusters the MAGs by similarity in counts between the different CAZyme domains, while the dendrogram above clusters CAZymes based on similarity in counts between the different MAGs. (B) Relative abundance of MAGs in the Alphaproteobacteria across host crab genera with a mean relative abundance >0.1% are displayed as a barplot. Bars are split by taxonomic genus, colored by predicted order and error bars represent standard error.

(A)

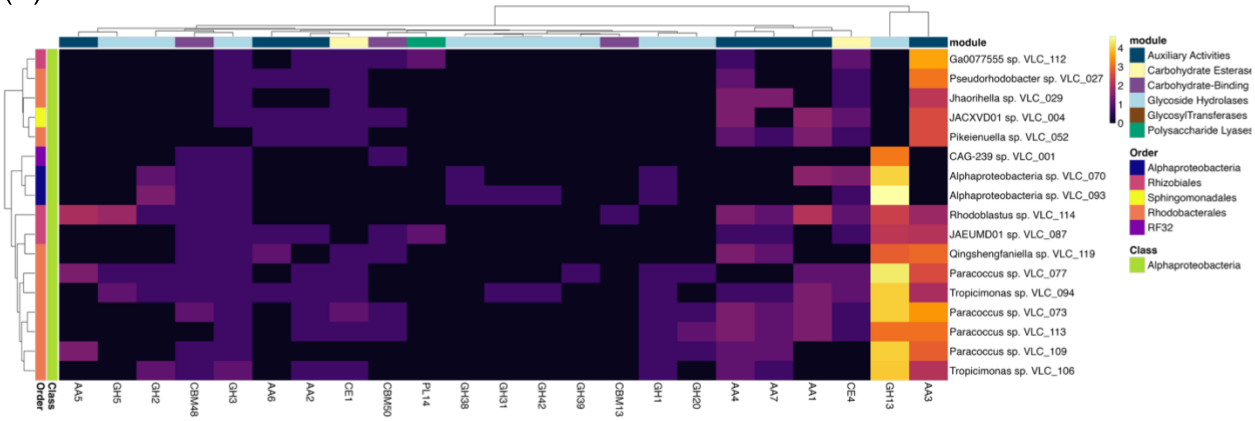


(B)

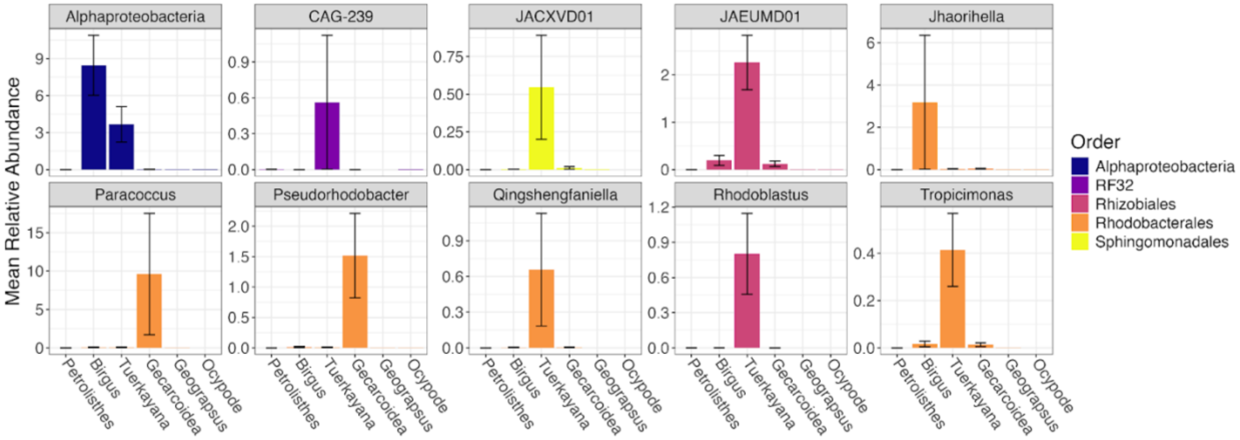

**Figure S9. CAZyme domain abundance across Bacilli.** (A) DRAM was used to annotate CAZyme domains in the proteome of MAGs. A heatmap displaying the log-transformed counts of CAZyme domains with potential lignocellulose-related functions for members of the Bacilli. Specific CAZyme modules are denoted across the top, while MAG taxonomic order and class information are denoted on the left. Further, the dendrogram to the left of the heatmap clusters the MAGs by similarity in counts between the different CAZyme domains, while the dendrogram above clusters CAZymes based on similarity in counts between the different MAGs. (B) Relative abundance of MAGs in the Bacilli across host crab genera with a mean relative abundance >0.1% are displayed as a barplot. Bars are split by taxonomic genus, colored by predicted order and error bars represent standard error.

(A)

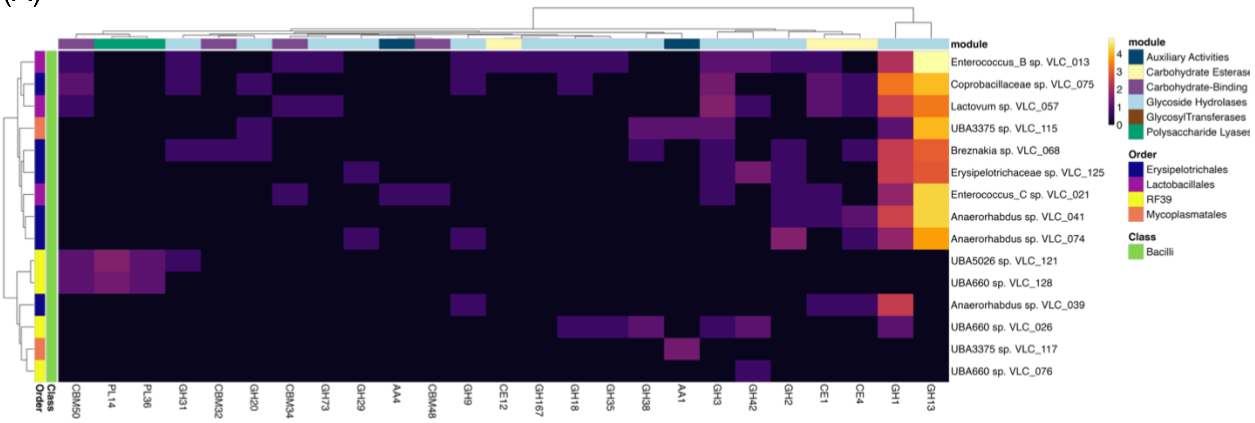


(B)

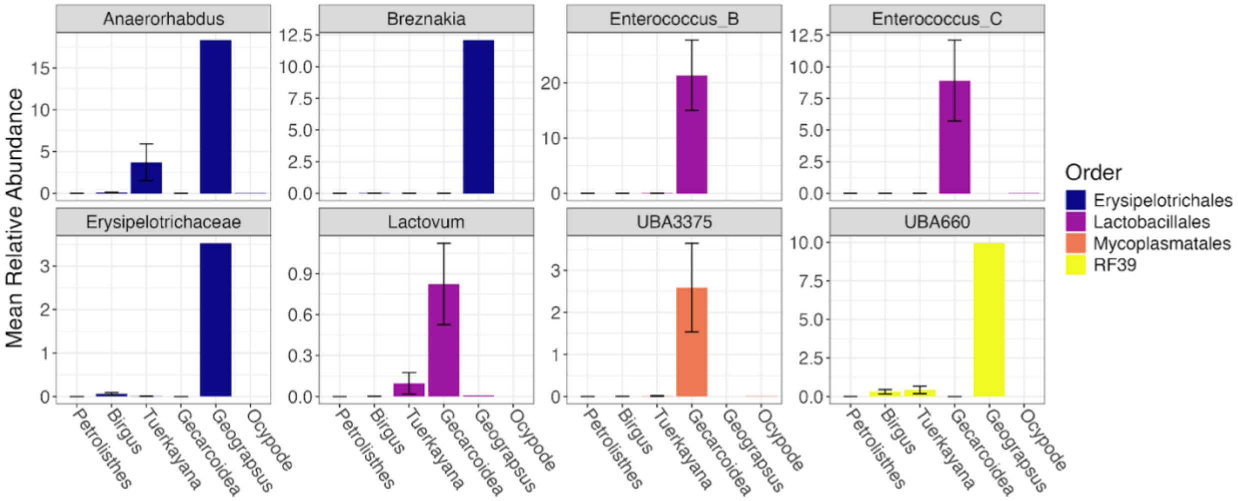

**Table S1. Taxonomic placement of MAGs by GTDB-Tk.** Full taxonomic assignment of each MAG from GTDB-Tk, as well as the calculated relative evolutionary divergence (RED) value of phylogenetic novelty. In cases where topological placement in the GTDB phylogeny and ANI result in congruent species assignments, GTDB-Tk does not report RED values (i.e., these values are reported as N/A).

| Bin ID | Phylum | Class | Order | Family | Genus | RED |
|---|---|---|---|---|---|---|
| VLC_001 | Proteobacteria | Alphaproteobacteria | RF32 | CAG-239 | | 0.75 |
| VLC_002 | Proteobacteria | Gammaproteobacteria | Burkholderiales | Burkholderiaceae | *Ottowia* | 0.95 |
| VLC_003 | Actinobacteriota | Actinomycetia | Actinomycetales | Demequinaceae | *Demequina* | 0.96 |
| VLC_004 | Proteobacteria | Alphaproteobacteria | Sphingomonadales | Sphingomonadaceae | *JACXVD01* | 0.98 |
| VLC_005 | Bacteroidota | Bacteroidia | Chitinophagales | Chitinophagaceae | *Niabella* | 0.98 |
| VLC_006 | Proteobacteria | Gammaproteobacteria | Burkholderiales | Rhodocyclaceae | *Denitromonas* | 0.99 |
| VLC_007 | Proteobacteria | Gammaproteobacteria | Burkholderiales | Burkholderiaceae | *Brachymonas* | 0.99 |
| VLC_008 | Campylobacterota | Campylobacteria | Campylobacterales | Arcobacteraceae | *Malaciobacter* | N/A |
| VLC_009 | Firmicutes_C | Negativicutes | Acidaminococcales | Acidaminococcaceae | *Phascolarctobacterium* | N/A |
| VLC_010 | Bacteroidota | Bacteroidia | Bacteroidales | P3 | *UBA10566* | N/A |
| VLC_011 | Fusobacteriota | Fusobacteriia | Fusobacteriales | Fusobacteriaceae | *Fusobacterium_A* | N/A |
| VLC_012 | Bacteroidota | Bacteroidia | Bacteroidales | Tannerellaceae | *Parabacteroides* | N/A |
| VLC_013 | Firmicutes | Bacilli | Lactobacillales | Enterococcaceae | *Enterococcus_B* | 0.90 |
| VLC_014 | Bacteroidota | Bacteroidia | Flavobacteriales | Flavobacteriaceae | *Flavobacteriaceae* | 0.96 |

| | | | | | | |
|---|---|---|---|---|---|---|
| VLC_015 | Bacteroidota | Bacteroidia | Bacteroidales | Bacteroidaceae | *Bacteroides* | N/A |
| VLC_016 | Bacteroidota | Bacteroidia | Flavobacteriales | Flavobacteriaceae | *Xanthomarina* | N/A |
| VLC_017 | Proteobacteria | Gammaproteobacteria | Enterobacterales | Vibrionaceae | *Vibrio* | 0.97 |
| VLC_018 | Bacteroidota | Bacteroidia | Bacteroidales | Bacteroidaceae | *Bacteroides* | 0.97 |
| VLC_019 | Bacteroidota | Bacteroidia | Bacteroidales | Prolixibacteraceae | *Draconibacterium* | 0.98 |
| VLC_020 | Proteobacteria | Gammaproteobacteria | Enterobacterales | Vibrionaceae | *Vibrio* | N/A |
| VLC_021 | Firmicutes | Bacilli | Lactobacillales | Enterococcaceae | *Enterococcus_C* | 0.90 |
| VLC_022 | Proteobacteria | Gammaproteobacteria | Enterobacterales | Enterobacteriaceae | *Citrobacter* | N/A |
| VLC_023 | Proteobacteria | Gammaproteobacteria | Enterobacterales | Shewanellaceae | *Shewanella* | 1.00 |
| VLC_024 | Proteobacteria | Gammaproteobacteria | Enterobacterales | Enterobacteriaceae | *Limnobaculum* | N/A |
| VLC_025 | Bacteroidota | Bacteroidia | Flavobacteriales | Flavobacteriaceae | *Aestuariibaculum* | 0.97 |
| VLC_026 | Firmicutes | Bacilli | RF39 | UBA660 | | 0.73 |
| VLC_027 | Proteobacteria | Alphaproteobacteria | Rhodobacterales | Rhodobacteraceae | *Pseudorhodobacter* | 0.97 |
| VLC_028 | Bacteroidota | Bacteroidia | Flavobacteriales | UJ101 | *UJ101* | 0.95 |
| VLC_029 | Proteobacteria | Alphaproteobacteria | Rhodobacterales | Rhodobacteraceae | *Jhaorihella* | 0.97 |
| VLC_030 | Bacteroidota | Bacteroidia | Bacteroidales | Prolixibacteraceae | *Mangrovibacterium* | 0.92 |
| VLC_031 | Actinobacteriota | Actinomycetia | Mycobacteriales | Antricoccaceae | *Cumulibacter* | 1.00 |

| | | | | | | |
|---|---|---|---|---|---|---|
| VLC_032 | Bacteroidota | Bacteroidia | Bacteroidales | Dysgonomonadaceae | *Dysgonomonas* | 0.97 |
| VLC_033 | Actinobacteriota | Actinomycetia | Mycobacteriales | Mycobacteriaceae | *Mycobacterium* | 0.99 |
| VLC_034 | Firmicutes_A | Clostridia | Lachnospirales | Lachnospiraceae | | 0.88 |
| VLC_035 | Bacteroidota | Bacteroidia | Bacteroidales | P3 | *UBA10566* | 0.92 |
| VLC_036 | Bacteroidota | Bacteroidia | Bacteroidales | | | 0.74 |
| VLC_037 | Firmicutes_A | Clostridia | Oscillospirales | Ruminococcaceae | *Fimivivens* | 0.94 |
| VLC_038 | Proteobacteria | Gammaproteobacteria | Xanthomonadales | Marinicellaceae | *Marinicella* | N/A |
| VLC_039 | Firmicutes | Bacilli | Erysipelotrichales | Erysipelotrichaceae | *Anaerorhabdus* | 0.90 |
| VLC_040 | Actinobacteriota | Actinomycetia | Actinomycetales | Dermatophilaceae | *Tetrasphaera* | 0.89 |
| VLC_041 | Firmicutes | Bacilli | Erysipelotrichales | Erysipelotrichaceae | *Anaerorhabdus* | 0.94 |
| VLC_042 | Firmicutes_A | Clostridia | Peptostreptococcales | Anaerovoracaceae | *Aminipila* | 0.90 |
| VLC_043 | Proteobacteria | Gammaproteobacteria | Pseudomonadales | Halieaceae | *Parahaliea* | 0.95 |
| VLC_044 | Firmicutes_A | Clostridia | Lachnospirales | Lachnospiraceae | | 0.89 |
| VLC_045 | Actinobacteriota | Actinomycetia | Actinomycetales | GCA-002748415 | | 0.76 |
| VLC_046 | Firmicutes_A | Clostridia | Lachnospirales | | | 0.70 |
| VLC_047 | Proteobacteria | Gammaproteobacteria | Enterobacterales | Shewanellaceae | *Shewanella* | N/A |
| VLC_048 | Actinobacteriota | Acidimicrobiia | Acidimicrobiales | SZUA-35 | *CADEDH01* | 0.88 |

| | | | | | | |
|---|---|---|---|---|---|---|
| VLC_049 | Patescibacteria | Saccharimonadia | Saccharimonadales | Nanosyncoccaceae | *UBA2834* | 0.91 |
| VLC_050 | Bacteroidota | Bacteroidia | Bacteroidales | Paludibacteraceae | *UPXZ01* | 0.95 |
| VLC_051 | Bacteroidota | Bacteroidia | Bacteroidales | | | 0.74 |
| VLC_052 | Proteobacteria | Alphaproteobacteria | Rhodobacterales | Rhodobacteraceae | *Pikeienuella* | 0.94 |
| VLC_053 | Firmicutes_A | Clostridia | Lachnospirales | | | 0.69 |
| VLC_054 | Bacteroidota | Bacteroidia | Bacteroidales | Dysgonomonadaceae | *Dysgonomonas* | 0.95 |
| VLC_055 | Bacteroidota | Bacteroidia | Bacteroidales | Prolixibacteraceae | *Mangrovibacterium* | 0.92 |
| VLC_056 | Campylobacterota | Campylobacteria | Campylobacterales | Campylobacteraceae | | 0.78 |
| VLC_057 | Firmicutes | Bacilli | Lactobacillales | Streptococcaceae | *Lactovum* | 0.96 |
| VLC_058 | Planctomycetota | Planctomycetia | Planctomycetales | Planctomycetaceae | | 0.81 |
| VLC_059 | Desulfobacterota | Desulfobulbia | Desulfobulbales | Desulfocapsaceae | *Desulfopila* | 0.95 |
| VLC_060 | Proteobacteria | Gammaproteobacteria | Pseudomonadales | Pseudomonadaceae | *Pseudomonas_A* | 1.00 |
| VLC_061 | Firmicutes_A | Clostridia | Lachnospirales | UBA5962 | | 0.74 |
| VLC_062 | Proteobacteria | Gammaproteobacteria | Enterobacterales | Vibrionaceae | *Vibrio* | 1.00 |
| VLC_063 | Actinobacteriota | Actinomycetia | Propionibacteriales | Propionibacteriaceae | *Brevilactibacter* | 0.96 |
| VLC_064 | Actinobacteriota | Actinomycetia | Propionibacteriales | Propionibacteriaceae | *Brevilactibacter* | 0.96 |
| VLC_065 | Acidobacteriota | Blastocatellia | Pyrinomonadales | Pyrinomonadaceae | *QQUY01* | 0.92 |

| | | | | | | |
|---|---|---|---|---|---|---|
| VLC_066 | Firmicutes_A | Clostridia | Oscillospirales | Ruminococcaceae | | 0.85 |
| VLC_067 | Firmicutes_A | Clostridia | Oscillospirales | Ruminococcaceae | *JAAYCI01* | 0.89 |
| VLC_068 | Firmicutes | Bacilli | Erysipelotrichales | Erysipelotrichaceae | *Breznakia* | 0.90 |
| VLC_069 | Firmicutes_A | Clostridia | Lachnospirales | Lachnospiraceae | *Suipraeoptans* | 0.95 |
| VLC_070 | Proteobacteria | Alphaproteobacteria | | | | 0.54 |
| VLC_071 | Actinobacteriota | Actinomycetia | Actinomycetales | Microbacteriaceae | *Leucobacter* | 0.97 |
| VLC_072 | Verrucomicrobiota | Verrucomicrobiae | Chthoniobacterales | Xiphinematobacteraceae | | 0.75 |
| VLC_073 | Proteobacteria | Alphaproteobacteria | Rhodobacterales | Rhodobacteraceae | *Paracoccus* | 0.98 |
| VLC_074 | Firmicutes | Bacilli | Erysipelotrichales | Erysipelotrichaceae | *Anaerorhabdus* | 0.89 |
| VLC_075 | Firmicutes | Bacilli | Erysipelotrichales | Coprobacillaceae | | 0.91 |
| VLC_076 | Firmicutes | Bacilli | RF39 | UBA660 | | 0.73 |
| VLC_077 | Proteobacteria | Alphaproteobacteria | Rhodobacterales | Rhodobacteraceae | *Paracoccus* | 0.98 |
| VLC_078 | Actinobacteriota | Actinomycetia | Propionibacteriales | Propionibacteriaceae | *Arachnia* | 0.94 |
| VLC_079 | Desulfobacterota_I | Desulfovibrionia | Desulfovibrionales | Desulfovibrionaceae | *Oleidesulfovibrio* | 0.93 |
| VLC_080 | Bacteroidota | Bacteroidia | Bacteroidales | Paludibacteraceae | *UPXZ01* | 0.94 |
| VLC_081 | Bacteroidota | Bacteroidia | Bacteroidales | Prolixibacteraceae | *Mangrovibacterium* | 0.92 |
| VLC_082 | Actinobacteriota | Actinomycetia | Actinomycetales | Beutenbergiaceae | | 0.91 |

| | | | | | | |
|---|---|---|---|---|---|---|
| VLC_083 | Bacteroidota | Bacteroidia | Bacteroidales | UBA7332 | | 0.86 |
| VLC_084 | Bacteroidota | Bacteroidia | Bacteroidales | Dysgonomonadaceae | *Dysgonomonas* | 0.95 |
| VLC_085 | Bacteroidota | Bacteroidia | Bacteroidales | Bacteroidaceae | *Bacteroides* | N/A |
| VLC_086 | Actinobacteriota | Actinomycetia | Actinomycetales | Microbacteriaceae | *Microbacterium* | 0.96 |
| VLC_087 | Proteobacteria | Alphaproteobacteria | Rhizobiales | Hyphomicrobiaceae | *JAEUMD01* | 0.93 |
| VLC_088 | Bacteroidota | Bacteroidia | Bacteroidales | Dysgonomonadaceae | *Dysgonomonas* | 0.97 |
| VLC_089 | Proteobacteria | Gammaproteobacteria | Burkholderiales | Rhodocyclaceae | *Azonexus* | N/A |
| VLC_090 | Bacteroidota | Bacteroidia | Bacteroidales | Dysgonomonadaceae | *Dysgonomonas* | N/A |
| VLC_091 | Bacteroidota | Bacteroidia | Bacteroidales | Bacteroidaceae | *Bacteroides* | N/A |
| VLC_092 | Firmicutes_A | Clostridia | Lachnospirales | Anaerotignaceae | *Anaerotignum* | 0.95 |
| VLC_093 | Proteobacteria | Alphaproteobacteria | | | | 0.53 |
| VLC_094 | Proteobacteria | Alphaproteobacteria | Rhodobacterales | Rhodobacteraceae | *Tropicimonas* | 0.98 |
| VLC_095 | Bacteroidota | Bacteroidia | Bacteroidales | Bacteroidaceae | *Bacteroides* | N/A |
| VLC_096 | Firmicutes_A | Clostridia | Lachnospirales | Lachnospiraceae | *Anaerocolumna* | 0.93 |
| VLC_097 | Bacteroidota | Bacteroidia | Bacteroidales | Prolixibacteraceae | *Mangrovibacterium* | 0.93 |
| VLC_098 | Firmicutes_A | Clostridia | Oscillospirales | Ruminococcaceae | *JAAYCI01* | 0.88 |
| VLC_099 | Chlamydiota | Chlamydiia | Chlamydiales | Parachlamydiaceae | | 0.85 |

| | | | | | | |
|---|---|---|---|---|---|---|
| VLC_100 | Desulfobacterota_I | Desulfovibrionia | Desulfovibrionales | Desulfovibrionaceae | *Halodesulfovibrio* | 0.98 |
| VLC_101 | Bacteroidota | Bacteroidia | Bacteroidales | | | 0.74 |
| VLC_102 | Actinobacteriota | Acidimicrobiia | Acidimicrobiales | Microtrichaceae | | 0.86 |
| VLC_103 | Actinobacteriota | Acidimicrobiia | Acidimicrobiales | Ilumatobacteraceae | *Desertimonas* | 0.88 |
| VLC_104 | Bacteroidota | Bacteroidia | Flavobacteriales | Weeksellaceae | *Moheibacter* | 0.93 |
| VLC_105 | Actinobacteriota | Actinomycetia | Actinomycetales | Microbacteriaceae | *Microbacterium* | 0.94 |
| VLC_106 | Proteobacteria | Alphaproteobacteria | Rhodobacterales | Rhodobacteraceae | *Tropicimonas* | 0.96 |
| VLC_107 | Verrucomicrobiota | Verrucomicrobiae | Chthoniobacterales | Xiphinematobacteraceae | | 0.75 |
| VLC_108 | Bacteroidota | Bacteroidia | Bacteroidales | Prolixibacteraceae | *Mangrovibacterium* | 0.92 |
| VLC_109 | Proteobacteria | Alphaproteobacteria | Rhodobacterales | Rhodobacteraceae | *Paracoccus* | 0.99 |
| VLC_110 | Proteobacteria | Gammaproteobacteria | Burkholderiales | Burkholderiaceae | *Castellaniella* | 0.99 |
| VLC_111 | Actinobacteriota | Actinomycetia | Actinomycetales | Demequinaceae | *Demequina* | 0.99 |
| VLC_112 | Proteobacteria | Alphaproteobacteria | Rhizobiales | Hyphomicrobiaceae | *Ga0077555* | 0.96 |
| VLC_113 | Proteobacteria | Alphaproteobacteria | Rhodobacterales | Rhodobacteraceae | *Paracoccus* | 0.99 |
| VLC_114 | Proteobacteria | Alphaproteobacteria | Rhizobiales | Beijerinckiaceae | *Rhodoblastus* | 0.92 |
| VLC_115 | Firmicutes | Bacilli | Mycoplasmatales | UBA3375 | | 0.84 |
| VLC_116 | Bacteroidota | Bacteroidia | Flavobacteriales | Flavobacteriaceae | *Tenacibaculum* | 0.98 |

| | | | | | | |
|---|---|---|---|---|---|---|
| VLC_117 | Firmicutes | Bacilli | Mycoplasmatales | UBA3375 | | 0.84 |
| VLC_118 | Bacteroidota | Bacteroidia | Flavobacteriales | UJ101 | *UJ101* | 0.97 |
| VLC_119 | Proteobacteria | Alphaproteobacteria | Rhodobacterales | Rhodobacteraceae | *Qingshengfaniella* | 0.95 |
| VLC_120 | Proteobacteria | Gammaproteobacteria | Pseudomonadales | Halieaceae | *Parahaliea* | 0.95 |
| VLC_121 | Firmicutes | Bacilli | RF39 | UBA660 | *UBA5026* | 0.93 |
| VLC_122 | Actinobacteriota | Actinomycetia | Actinomycetales | Beutenbergiaceae | | 0.92 |
| VLC_123 | Actinobacteriota | Acidimicrobiia | Acidimicrobiales | SZUA-35 | *CADEDH01* | 0.88 |
| VLC_124 | Firmicutes_A | Clostridia | Lachnospirales | Lachnospiraceae | *Lacrimispora* | N/A |
| VLC_125 | Firmicutes | Bacilli | Erysipelotrichales | Erysipelotrichaceae | | 0.78 |
| VLC_126 | Bacteroidota | Bacteroidia | Bacteroidales | Prolixibacteraceae | *Draconibacterium* | 1.00 |
| VLC_127 | Bacteroidota | Bacteroidia | Bacteroidales | Tannerellaceae | *Macellibacteroides* | N/A |
| VLC_128 | Firmicutes | Bacilli | RF39 | UBA660 | | 0.78 |
| VLC_129 | Proteobacteria | Gammaproteobacteria | Enterobacterales | Enterobacteriaceae | *Kluyvera* | 1.00 |

**Table S2. Beta diversity analysis.** Results of overall permutational multivariate analysis of variances (PERMANOVAs) using adonis2, *post-hoc* pairwise PERMANOVA comparisons using pairwise.adonis, and multivariate homogeneity of group dispersions using betadisper to assess gut microbiome beta diversity across crab genera, diets and terrestrial grades.

**Table S3. *Post-hoc* Tukey's Honest Significant Difference (HSD) test of dispersion.** Results of Tukey's HSD for pairwise comparisons of dispersion among crab genera, diets and terrestrial grades.

**Table S4. Kruskal-Wallis (KW) tests of relative abundance.** Results of KW tests evaluating significant differences in the relative abundance of microbial taxa among crab genera, diets and terrestrial grades.

**Table S5. *Post-hoc* Dunn tests of relative abundance.** *Post-hoc* Dunn tests for pairwise comparisons of microbial relative abundance following significant Kruskal-Wallis tests, detailing specific differences among crab genera, diets and terrestrial grades.

**Table S6. Random forest classification results.** Results of random forest classification predicting crab diet type based on METABOLIC complex carbon degradation pathway gene counts, including the confusion matrix summarizing model performance and ranked feature importance scores for diet classification.
